# TRAK adaptors coordinate the recruitment and activation of dynein and kinesin to control mitochondrial transport

**DOI:** 10.1101/2021.07.30.454553

**Authors:** John T. Canty, Andrew Hensley, Ahmet Yildiz

**Author notes:** Correspondence (Yildiz, A.).

## Abstract

In neurons, mitochondria are transported to distal regions for supplying energy and buffer Ca^2+^. Mitochondrial transport is mediated by Miro and TRAK adaptors that recruit kinesin and dynein-dynactin. To understand how mitochondria are transported by these opposing motors and stalled at regions with elevated Ca^2+^, we reconstituted the mitochondrial transport machinery in vitro. We show that the coiled-coil domain of TRAK activates dynein-dynactin motility, but kinesin requires an additional factor to efficiently transport Miro/TRAK. Unexpectedly, TRAK adaptors that recruit both motors move towards kinesin’s direction, whereas kinesin is excluded from binding TRAK transported by dynein-dynactin. The assembly and motility of the transport machinery are not affected by Ca^2+^. Instead, the mitochondrial docking protein syntaphilin is sufficient to oppose the forces generated by kinesin and stall the motility. Our results provide mechanistic insight into how mitochondria are transported by the coordinated action of motors and statically anchored to regions with high neuronal activity.

---

Mitochondria are cellular power plants that generate most of the ATP needed for many biochemical reactions and have a high capacity to buffer cytosolic Ca^2+^. In neurons, mitochondria are distributed to distal areas where energy and Ca^2+^ buffering are in high demand, such as synapses and axonal branches^8^. Mitochondrial transport is essential for axonal growth and branching, maintaining action potentials, and supporting the transport of synaptic transmission^1^. Aged and dysfunctional mitochondria need to be transported back to the cell body for degradation^9^. Defects in mitochondrial transport are associated with a variety of neurodegenerative diseases^2^

How mitochondrial trafficking is properly regulated to satisfy local energy requirements is not well understood. The complex transport properties of mitochondria are driven by receptor molecules, adaptors, and motor proteins. Early genetic screens identified the mitochondrial Rho GTPase Miro1^10^ at the center of mitochondrial cellular function. Miro1 is composed of two GTPase domains, two EF-hands that bind Ca^2+^, and the C-terminal transmembrane domain (TM). Miro1 is localized to the outer mitochondrial membrane through its TM^15,16^, and is coupled to C-terminus of the trafficking of kinesin-binding (TRAK) adaptors ^11^. The N-terminal coiled-coil domain of TRAK1 and TRAK2 can separately bind to the kinesin-1 heavy chain and dynein-dynactin^14,15^ (hereafter, kinesin and dynein), which transport mitochondria towards the plus- and minus-ends of microtubules, respectively^12–14^. Co-immunoprecipitation studies in neurons showed that TRAK1 recruits both dynein and kinesin whereas TRAK2 primarily interacts with dynein^14^. TRAK1 was found to be enriched in the axons of cultured neurons, while TRAK2 was found mainly in dendrites^14^, suggesting that these adaptors may have nonredundant roles in mitochondrial trafficking.

Live cell-imaging of cultured neurons has shown that mitochondria exhibit rapid anterograde and retrograde transport, interspersed with pausing and directional switching in axons and dendrites^3–7^. The complex transport properties of mitochondria are primarily driven by Miro1, TRAK adaptors, motors, and other associated factors, but little is known about the mechanism of the mitochondrial transport machinery. In vitro reconstitution studies showed that TRAK1 binds and activates kinesin motility^17^, but the association of TRAK with the dynein/dynactin transport machinery has not been demonstrated. It also remains unclear whether TRAK adaptors can simultaneously recruit both kinesin and dynein and coordinate their activity to control the bidirectional transport of mitochondria.

In mature neurons, only one-third of the mitochondria are transported, whereas two-thirds remain docked to microtubules and satisfy energy requirements and buffer Ca^2+^in regions with high neuronal activity^8^. Studies in cultured neurons demonstrated that mitochondrial transport stalls when local Ca^2+^ concentrations are elevated^8^, but the underlying mechanism remains controversial^8,16,18^. The ‘motor detachment’ model claims that Ca^2+^ binding to Miro1 decouples kinesin from TRAK^16^. In the ‘Miro-binding’ model, Ca^2+^ binding causes the motor domains of kinesin to detach from the microtubule (MT) and bind to Miro1^18^. In both cases, decoupling or inactivation of kinesin stalls the anterograde transport of mitochondria. However, elevated Ca^2+^ concentration also stops mitochondrial transport in the retrograde direction^19,20^. In addition, knockdown of Miro1 in neurons does not completely inhibit Ca^2+^-induced arrest of mitochondria^19,20^, suggesting that the Ca^2+^ mediated arrest may function independently of the transport machinery.

Recent studies in mouse models proposed an alternative model, in which a mitochondrial docking protein, syntaphilin (SNPH) anchors mitochondria to MTs in axons^21^. SNPH is recruited to mitochondria in response to sustained neuronal activity and elevated Ca^2+^ levels^21^. This model is supported by the observations that overexpression of SNPH completely abolishes mitochondrial transport whereas increasing the cytosolic Ca^2+^ fails to arrest mitochondrial transport in axons of SNPH knock-out neurons^21,22^. The “engine-switch and brake” model proposes that SNPH inhibits kinesin through direct molecular interaction^22^ and serves as a brake by anchoring mitochondria to MTs^21^. These models make specific predictions about how Ca^2+^ regulates transport and stalls mitochondria, but these predictions could not be directly tested due to the lack of a reconstituted system from purified components of the mitochondrial transport machinery.

In this study, we investigated the role of Miro and TRAK in the recruitment and activation of dynein and kinesin motility using *in vitro* reconstitution. We show that TRAK1 and TRAK2 activate dynein-dynactin motility. These adaptors also recruit but do not fully activate kinesin, which requires an additional factor for robust motility. TRAK1 or TRAK2 simultaneously recruits both dynein-dynactin and kinesin, but it coordinates the activity of opposing motors to avoid futile tug-of-war and determine the directionality of transport. We also observed that Miro1 stably interacts with kinesin/TRAK, but the motility of this complex is unaffected by excess Ca^2+^. However, static anchoring by SNPH is sufficient to stall kinesin motility. These results provide critical insight into how Miro/TRAK adaptors regulate the bidirectional motility of mitochondria.

## Results

### TRAK1 and TRAK2 are activating adaptors of dynein-dynactin

Recent studies have identified a family of coiled-coil adaptor proteins that activate dynein motility by recruiting one or two dynein motors to dynactin^23–25^. Sequence alignments with established dynein adaptors confirmed that TRAK1 and TRAK2 contain the CC1 box that binds the dynein light-intermediate chain (LIC)^26,27^ and the Spindly motif that interacts with the pointed-end of dynactin^28^ **(Fig. 1a,b)**. We first investigated whether human TRAK1/2 could activate mammalian dynein-dynactin for processive motility using single-molecule imaging *in vitro* **(Extended Data Fig. 1a)**. In the absence of TRAK, dynein and dynactin exhibited little to no motility, as previously shown^29–31^ **(Extended Data Fig. 1b)**. Remarkably, the N-terminal coiled coils of TRAK1/2 that contain both the CC1 box and the Spindly motif (TRAK1^1-400^ and TRAK2^1-400^) led to robust activation of dynein-dynactin motility towards the MT minus-end **(Fig. 1c-e, Extended Data Fig. 1b, and Supplementary Video 1)**. The velocities of dynein-dynactin-TRAK1^1-400^ (DDT_1_^1-400^), and -TRAK2^1-400^ (DDT_2_^1-400^) (810 ± 20 and 870 ± 20 nm s^−1^, mean ± s.e.m., respectively) were comparable to that of dynein-dynactin assembled with BicD adaptors in vitro^29,30,32,33^ and the retrograde transport speed of mitochondria (300 – 900 nm s^−1^) *in vivo*^3,14^ **(Extended Data Fig. 1c)**. In comparison, TRAK1/2 constructs that contain the CC1 box but lack the Spindly motif (TRAK1^1-360^ and TRAK2^1-360^) resulted in only occasional motility (**Fig. 1c,d)**, underscoring the importance of the Spindly motif in the activation of dynein-dynactin.

**Fig. 1.**
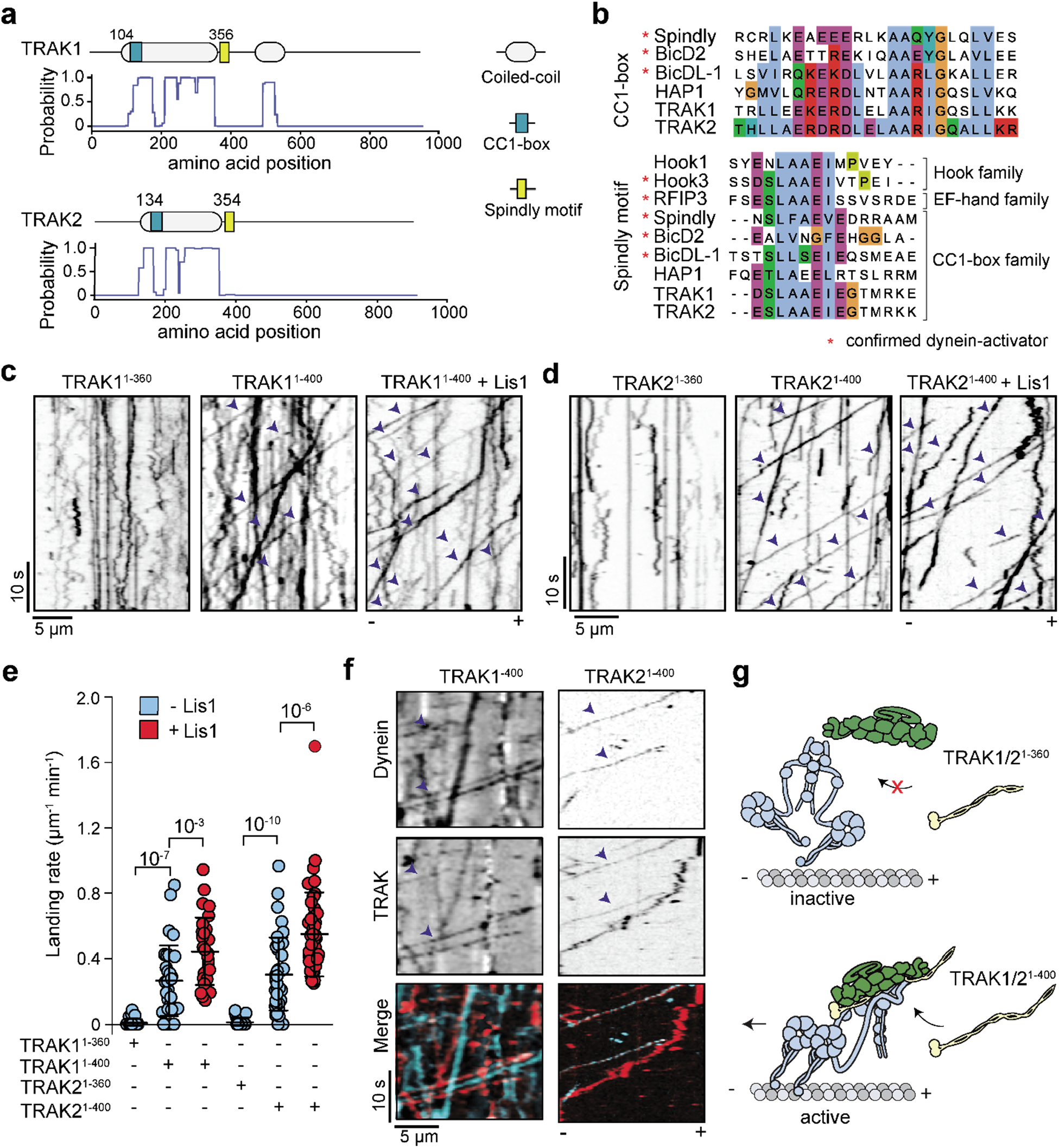
TRAK coiled-coil domains activate dynein-dynactin motility. **a**, Domain organization and coiled-coil prediction score of human TRAK1 and TRAK2. **b**, Sequence alignment shows the conserved CC1 box (top) and the Spindly motif (bottom) of TRAK1 and TRAK2 with other activating adaptors of human dynein-1. **c-d**, Representative kymographs of TRAK1 (**c**) and TRAK2 (**d**) constructs labeled with the LD655 dye in the presence of unlabeled dynein/dynactin (DD) and Lis1. Arrowheads highlight processive motility. **e**, The landing rate of motor complexes on MTs. The centerline and whiskers represent mean and s.d., respectively (*n* = 30, 32, 31, 31, 44, and 48 MTs from left to right). *P* values are calculated from a two-tailed *t*-test. **f**, Representative two-color kymograph of LD655-dynein and LD555-labeled TRAK constructs in the presence of unlabeled dynactin. Arrowheads represent TRAK-dynein colocalization. The processive motility of TRAK not colocalizing with dynein is due to less than 100% labeling efficiency of dynein. **g**, TRAK is an activating adaptor of dynein-dynactin. Activation of dynein motility requires both the CC1 box and the Spindly motif in the TRAK coiled-coil.

Multi-color tracking experiments visualized direct colocalization of motile dynein and TRAK adaptors **(Fig. 1f)**, indicating that TRAK needs to be part of the dynein-dynactin complex to sustain processive motility. The MT landing rate of active DDT_1_^1-400^ and DDT_2_^1-400^ complexes increased two-fold by the addition of 1 μM Lis1 (**Fig. 1e)**, a dynein regulatory protein that facilitates the assembly of active dynein-dynactin-adaptor complexes^34–36^. Collectively, our results showed that TRAK1 and TRAK2 are activating adaptors of dynein-dynactin for mitochondrial transport **(Fig. 1g)**.

### TRAK recruits but does not efficiently activate kinesin

Previous studies reported that TRAK binds to kinesin between residues 100-360 of its coiled-coil domain, and activates kinesin motility on MTs^14,17^. We tested this by measuring the landing rate and mobile fraction of full-length human kinesin-1 heavy chain (KIF5B) with and without TRAK adaptors. In the absence of TRAK, kinesin landed infrequently onto the MTs and had a low mobile fraction **(Fig. 2a,b)**, consistent with autoinhibition of this motor when not transporting a cargo^37^. In the presence of TRAK1^1-360^ and TRAK2^1-360^, kinesin motors co-localized with TRAK adaptors and moved at similar velocities to kinesin alone **(Fig. 2a,b, and Extended Data Fig. 2a-c)**, as previously reported^17^. We observed that kinesin co-localized more frequently with TRAK1^1-360^ than TRAK2^1-360^ **(Extended Data Fig. 2c)**. In contrast to a previous report^17^, the landing rate of kinesin increased only 1.5-fold in the presence of TRAK1^1-360^ and was unaffected by TRAK2^1-360^ **(Fig. 2b)**. Similar results were obtained when we used longer (TRAK1^1-400^ than TRAK2^1-400^) or full-length TRAK1^1-953^ and TRAK2^1-914^ constructs **(Extended Data Fig. 2d,e)**. Although the C-terminus of TRAK was reported to interact with MTs and increase the processivity of the kinesin-TRAK (KT) complex^17^, we did not observe MT binding of TRAK1^1-953^ and TRAK2^1-914^ without colocalizing with kinesin **(Extended Data Fig. 2f)**. These results show that TRAK coiled-coil domains stably bind, but do not fully activate kinesin motility on MTs.

**Fig. 2.**
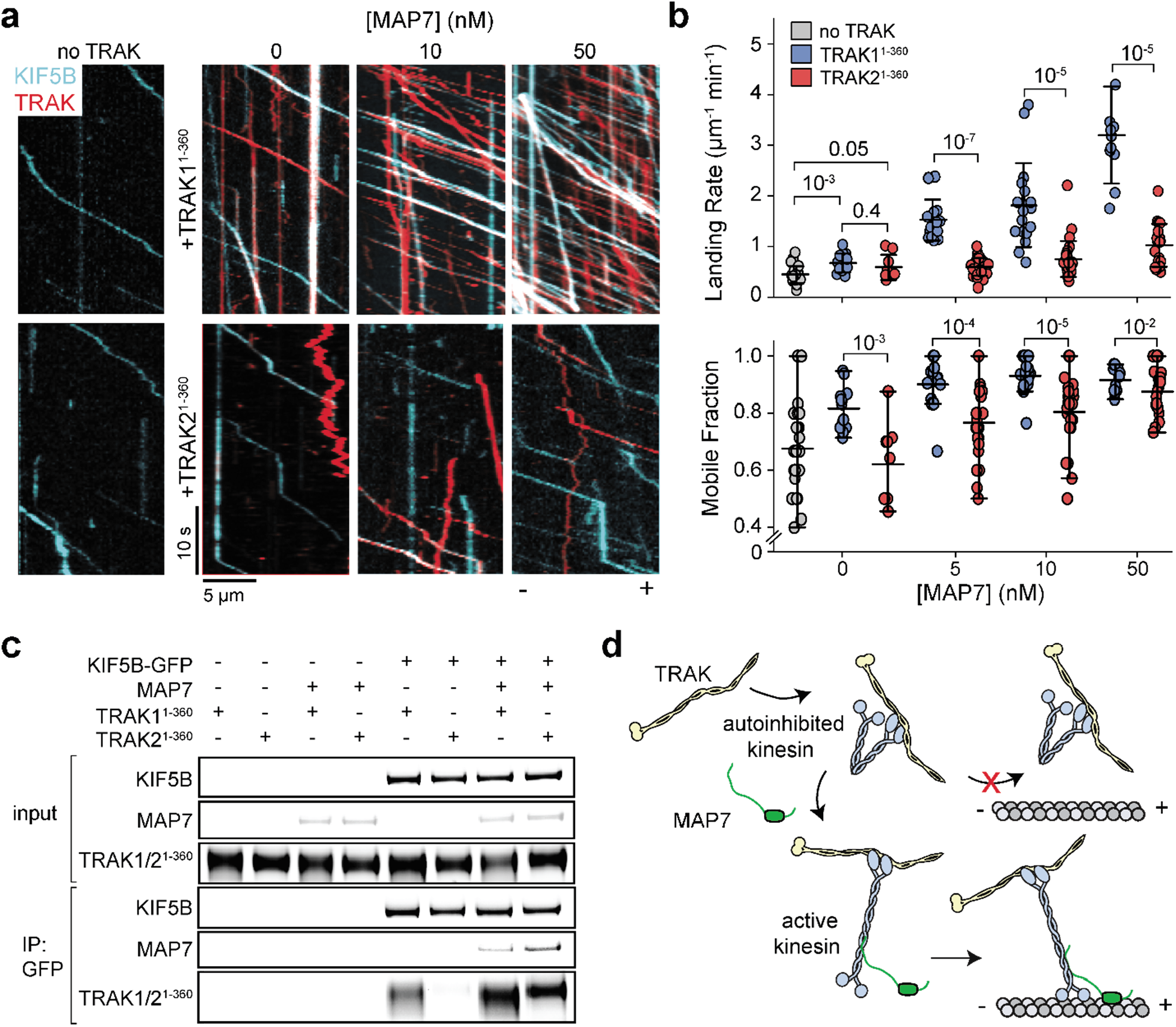
Kinesin recruits TRAK adaptors more efficiently following activation by MAP7. **a**, Representative two-color kymographs of kinesin (KIF5B) and TRAK constructs with increasing concentrations of MAP7. The processive motility of TRAK not colocalizing with kinesin is due to less than 100% labeling of kinesin. **b**, The landing rate and the mobile fraction of kinesin in the absence and presence of TRAK1^1-360^ or TRAK2^1-360^ under increasing MAP7 concentrations. The center line and whiskers represent the mean and s.d., respectively (*n* = 26, 13, 14, 18, 12, 9, 22, 29, and 19 MTs from left to right, three independent trials). P values are calculated from a twotailed t-test. **c**, In vitro immunoprecipitation (IP) of purified kinesin (KIF5B-GFP), TRAK1^1-360^, TRAK2^1-360^, in the presence or absence of MAP7. The proteins were eluted from anti-GFP beads. **d**, Schematics show that TRAK recruits kinesin, but activation of kinesin/TRAK motility requires an additional factor, such as MAP7.

We next asked whether TRAK adaptors more efficiently recruit kinesin when this motor is activated by additional factors, such as MAP7^38,39^. Consistent with previous reports^38,39^, the landing rate and mobile fraction of kinesin increased substantially under the increasing concentrations of MAP7 **(Fig. 2a,b and Supplementary Video 2)**. The landing rate of kinesin-/TRAK1^1-360^ (KT_1_^1-360^) complexes increased more than 7-fold, while kinesin-1/TRAK2^1-360^ (KT_2_^1-360^) increased only 2-fold by 50 nM MAP7 **(Figure 2b)**. Pull-down assays also showed that kinesin binds to TRAK1^1-360^, but not TRAK2^1-360^ without MAP7, but it binds to both adaptors in the presence of MAP7 **(Fig. 2c)**. Collectively, these results demonstrate that the coiled-coil domain of TRAK1, and to a lesser extent TRAK2, more efficiently recruits kinesin when the motor is rescued from autoinhibition **(Fig. 2d)**.

### Force generation of complexes formed with TRAK adaptors

To test how TRAK binding affects the force production of dynein and kinesin, we measured the stall forces of KT_1_^1-360^, DDT_1_^1-400^, and DDT2^1-400^ complexes using an optical trap **(Extended Data Fig. 3 a,b)**. To ensure that the forces measured corresponded to a fully assembled complex, we attached the beads directly to TRAK adaptors using a GFP-antibody linkage **(Fig. 3)**. Both DDT_1_^1-400^ and DDT_2_^1-400^ complexes stalled when subjected to 4.3 pN resistive forces **(Fig. 3)** and exhibited similar stall times before MT detachment **(Extended Data Fig. 3c,d)**. These forces are comparable to that of complexes that contain a single dynein motor and are lower than the complexes that contain two dyneins^23,33^, suggesting that TRAK primarily recruits one dynein to dynactin in our reconstitution conditions. KT_1_^1-360^ stalled at 5.96 ± 0.24 pN load **(Fig. 3)**, which closely matched to measured stall forces of constitutively active kinesin^40^, indicating that TRAK recruits a single kinesin motor^17^. We were unable to detect any bead motility in the presence of TRAK2^1-360^, consistent with the low affinity of kinesin for this construct. These results show that DDT and KT are active complexes that generate sufficient force to drive retrograde and anterograde motility.

**Fig. 3.**
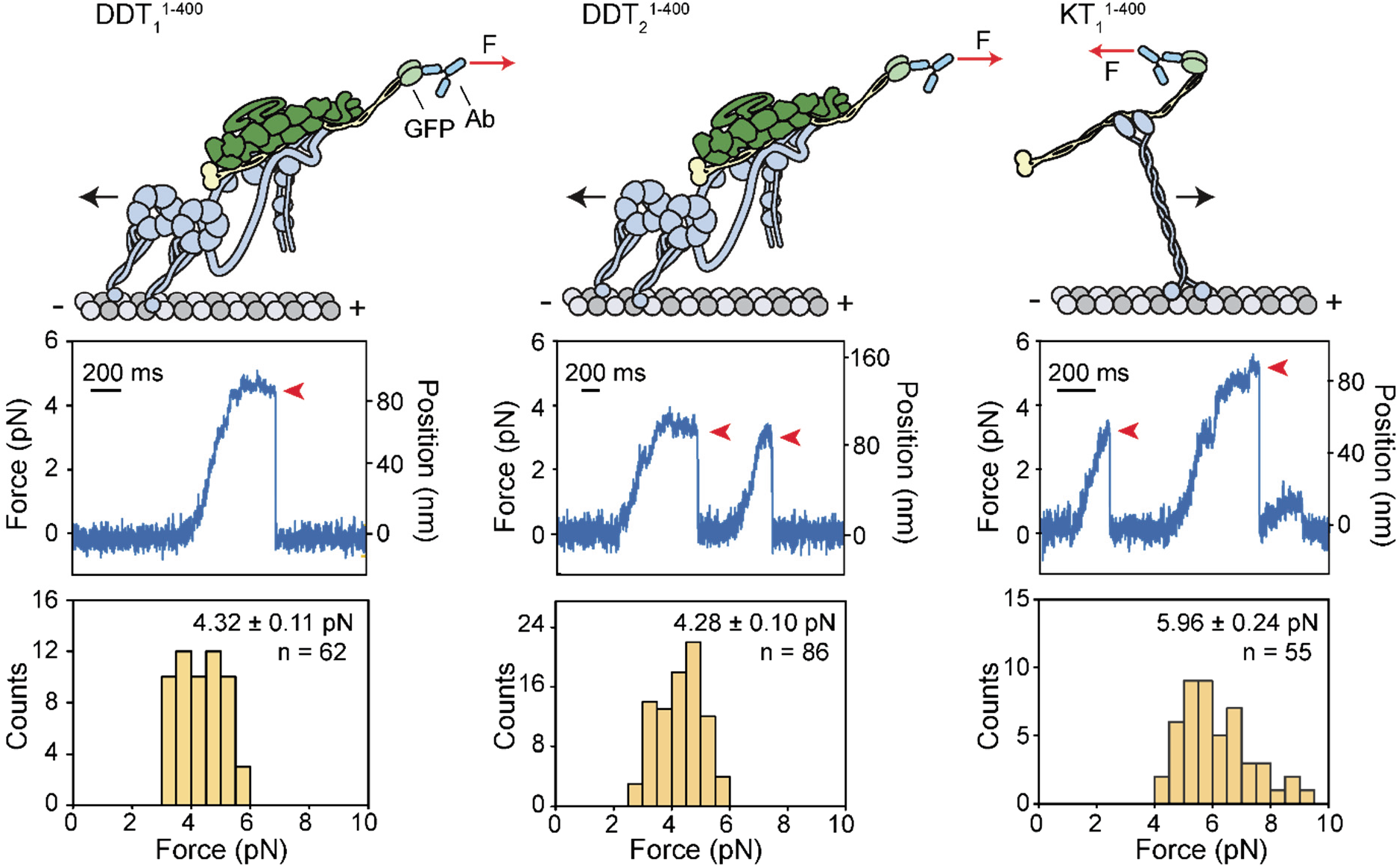
Force generation of dynein-dynactin or kinesin assembled with TRAK adaptors. (Top) Single motor complexes were pulled from the TRAK adaptor by an optically trapped (not shown) under force (F). (Middle) Representative traces of beads driven by a single complex in a fixed-trap assay. Red arrowheads represent the detachment of the motor from an MT after the stall. (Bottom) Stall forces (mean ± s.e.m.) of DDT_1_^1-400^ (*n* = 62 stalls from 15 beads in 7 independent experiments), DDT_2_^1-400^ (*n* = 86 stalls from 14 beads in 6 independent experiments) and KT_1_^1-400^ (*n* = 55 stalls from 14 beads in 6 independent experiments).

### TRAK adaptors simultaneously recruit dynein-dynactin and kinesin

A subset of activating adaptors has been shown to recruit both kinesin and dynein to form a bi-directional scaffold^41^. We performed three-color imaging of dynein-dynactin, kinesin, and either TRAK1^1-400^ or TRAK2^1-400^ to test whether dynein and kinesin can colocalize to the same TRAK adaptor. We observed dynein-dynactin/kinesin/TRAK1^1-400^ (DDKT_1_^1-400^) colocalizers moving along the MT **(Fig. 4a,b and Supplementary Video 3)**. The likelihood of detecting dynein and kinesin to simultaneously colocalize on TRAK2^1-400^ was substantially lower, presumably because kinesin has a low affinity for TRAK2 **(Fig. 4b)**. The analysis of these trajectories revealed that all DDKT_1_^1-400^ and DDKT_2_^1-400^ assemblies moved towards the MT plus-end at comparable velocities to KT_1_^1-400^ and KT_2_^1-400^ complexes in the same chamber **(Fig. 4b)**. We also noticed that DDKT complexes moved much faster than the case in which kinesin and dynein engage in a tug-of-war on an artificial DNA scaffold^32,33,43^. Collectively, these results indicate that dynein is not competing against kinesin-driven motility when both motors are recruited by TRAK^42^.

**Fig. 4.**
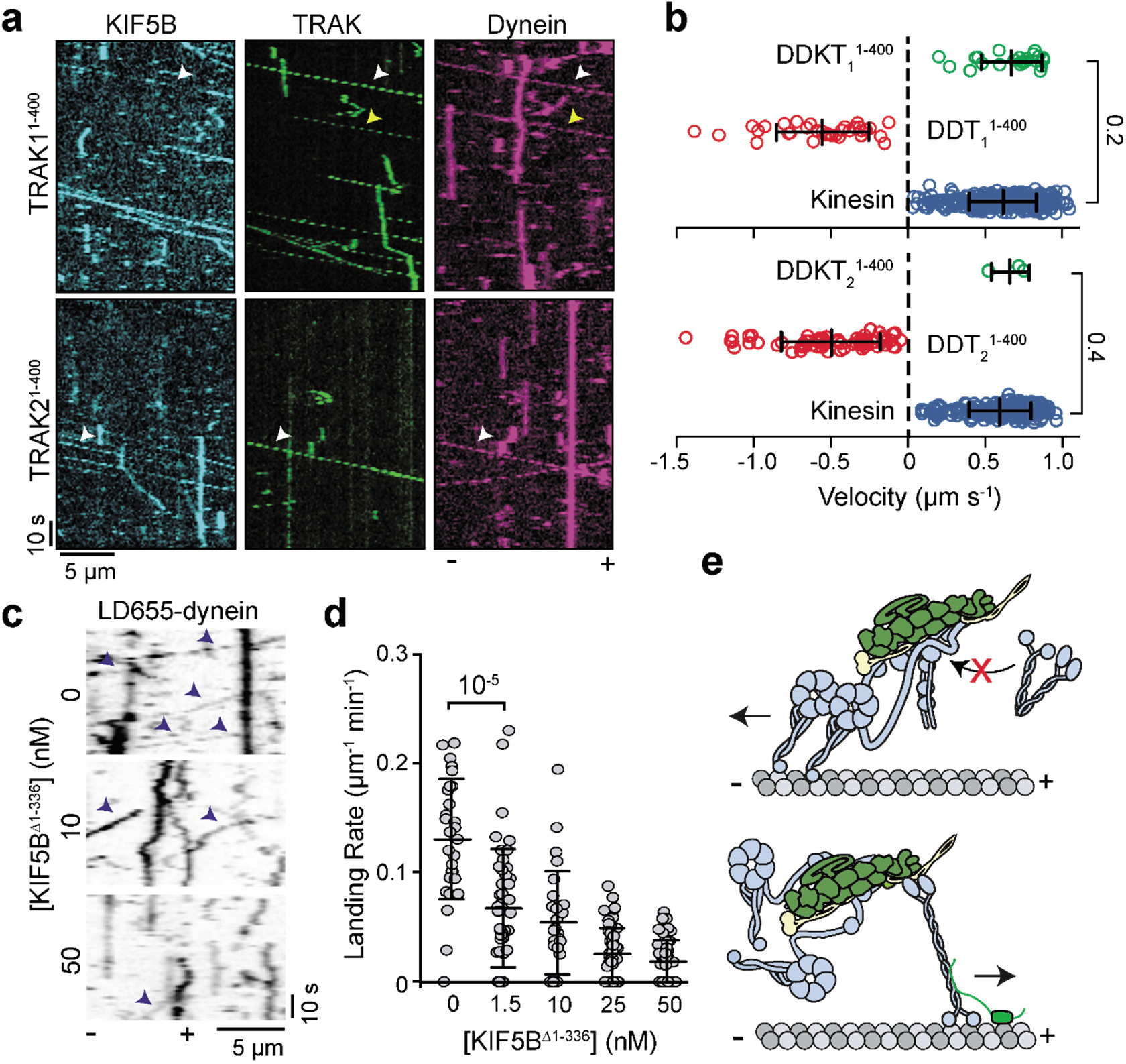
TRAK adaptors simultaneously recruit dynein-dynactin and kinesin. **a**, Sample kymographs of Alexa488-kinesin, LD555-TRAK1^1-400^ or TRAK2^1-400^, and LD655-dynein in the presence of 5 nM MAP7. White arrowheads show colocalization of dynein, kinesin, and TRAK. Yellow arrowheads highlight the plus-end-directed movement of dynein and TRAK by unlabeled kinesin. **b**, The velocity distribution of individual motor complexes and DDKT assemblies. Negative velocities correspond to minus-end-directed motility (*n* = 24, 37, 310, 3, 67, and 206 from top to bottom, with three independent experiments per condition). **c**, Sample kymographs of DDT1^1-400^ assemblies in the presence of 1 μM Lis1 and increasing concentrations of KIF5B^Δ1-336^. The motility of LD655-dyneins is highlighted with orange arrowheads. **d**, The landing rates of DDT_1_^1-400^ assemblies (0.5 nM) under increasing unlabeled KIF5B^Δ1-336^ concentrations (*n* = 28, 45, 27, 50, and 51 MTs from left to right). **e**, Schematic representation of motor coordination on TRAK. When kinesin transports TRAK, dynein can remain as an inactive passenger, but kinesin is excluded from minus-end directed DDT complexes. In (**b**) and (**d**), the center line and whiskers represent the mean and s.d., respectively. P-values are calculated from a two-tailed t-test.

We next investigated why DDKT complexes exclusively move towards the plus-end. We first tested whether dynein more efficiently competes against kinesin when its assembly with dynactin and TRAK is aided by Lis1^34–36^. The addition of 1 μM Lis1 had only a minor effect on the run frequency of complexes containing both motors, and no effect in their directional preference **(Extended Data Fig. 4)**, ruling out this possibility. DDKT may also be driven by higher force generation of kinesin compared to single dynein **(Fig. 3)**. To test this possibility, we replaced full-length kinesin with a construct that binds to TRAK but does not generate force and motility (KIF5B^Δ1-336^) and ask whether DDKT complexes assembled with KIF5B^Δ1-336^ move towards the minus end. However, the addition of excess KIF5B^Δ1-336^ resulted in more than a 5-fold reduction in DDT_1_^1-400^ motility **(Fig. 4c,d)** and we did not observe any KIF5B^Δ1-336^ colocalizing with processive DDT_1_^1-400^ complexes. These results indicate that when kinesin binds to TRAK, dynein/dynactin cannot form an active motor and is transported by kinesin towards the plus-end **(Fig. 4e)**.

### Miro1 forms a processive complex with kinesin and TRAK adaptors

We turned our attention to the association of Miro1 with the KT complex. Miro1 was shown to interact separately with TRAK adaptors and kinesin in immunoprecipitation assays^15,16,18^, but it remained unclear which of these interactions form a stable complex capable of plus-end directed transport. To address this question, we expressed a Miro1 construct lacking its C-terminal transmembrane domain (Miro1^1-592^, **Fig. 5a, Extended Data Fig. 5a)**. In the absence of TRAK adaptors, we observed colocalization of Miro1 to kinesin motors that walked along MTs **(Fig. 5b,c, Extended Data Fig. 5b,c, Supplementary Videos 4 and 5)**. Consistent with a previous report^16^, the addition of Ca^2+^ resulted in an 80% reduction of Miro1/kinesin colocalizers **(Fig. 5b,c)**, indicating that Miro1 dissociates from kinesin in a Ca^2+^ dependent manner. The introduction of either TRAK1^1-360^ or TRAK2^1-360^, which can bind to kinesin but not to Miro1, substantially reduced the number of motile Miro1-kinesin colocalizers in motility assays **(Fig. 5d,e, Extended Data Fig. 5d)** and disrupted Miro1-kinesin binding in co-immunoprecipitation assays **(Extended Data Fig. 5e).** Therefore, the N-terminal domain of TRAK1/2 competes with Miro1 for binding to the kinesin cargo binding domain.

**Fig. 5.**
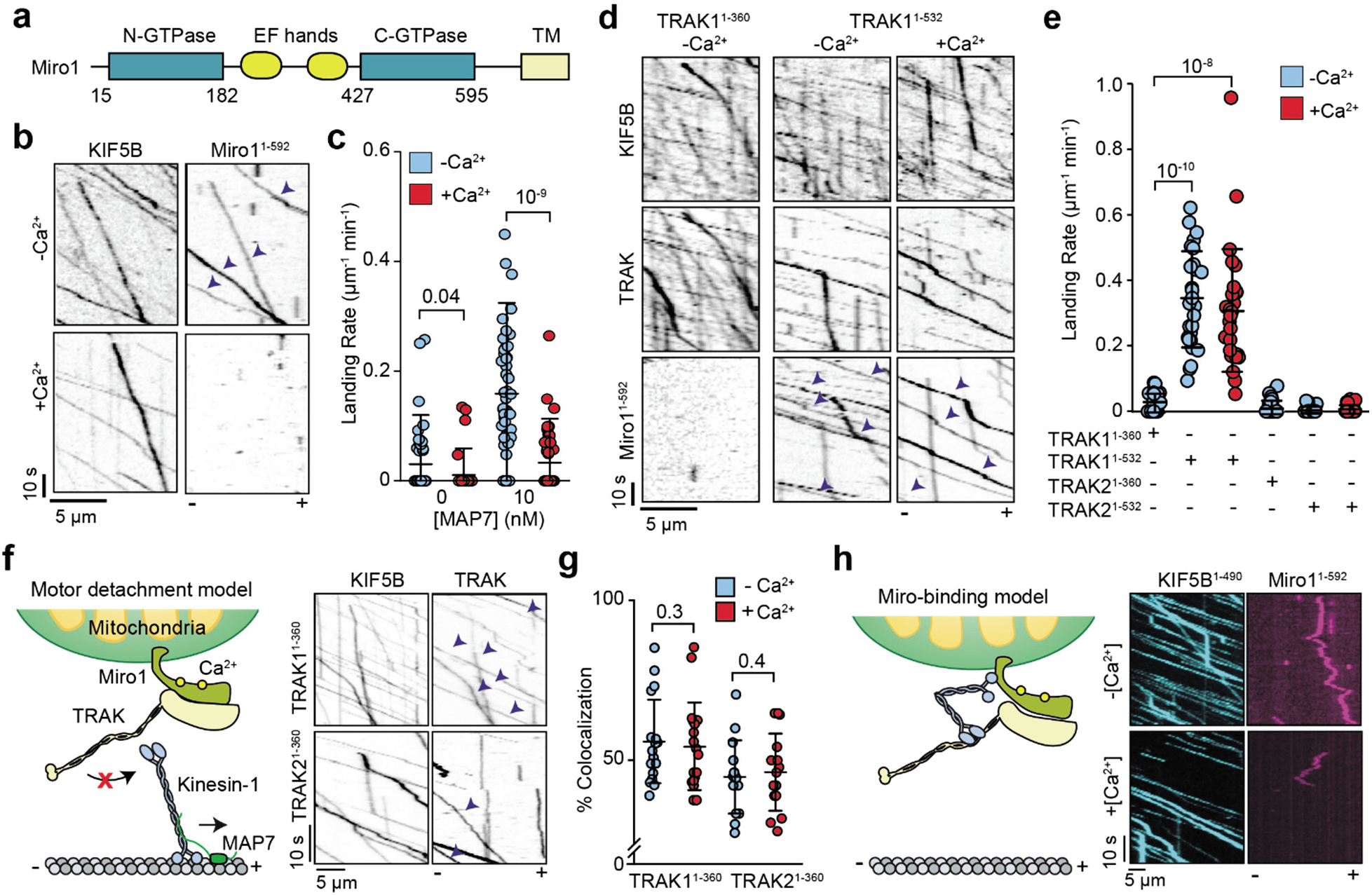
Miro1 forms a ternary complex with kinesin and TRAK1. **a**, Miro1 binds to GTP at its GTPase domains and Ca^2+^ ions at its EF-hands, and localizes to the mitochondrial outer membrane through its TM. **b**, Kymographs of LD655-kinesin (KIF5B) and LD555-Miro1^1-592^ in the presence and absence of 2 mM Ca^2+^. Assays were conducted in the presence of 10 nM MAP7 and the absence of TRAK adaptors. **c**, The landing rate of kinesin-Miro1 co-localizers (*n* = 47, 43, 45, and 50 MTs from left to right, three independent trials). **d**, Kymographs of Alexa488-kinesin, LD655-Miro1^1-592^, and LD555-labeled TRAK1 constructs. Assays were conducted in the presence of 10 nM MAP7. **e**, The landing rate of KTM co-localizers in the presence of different TRAK constructs (*n* = 27 MTs for each condition, three independent trials). **f**, (Left) The motor detachment model predicts that Ca^2+^ binding to Miro1 triggers dissociation of kinesin from TRAK. (Right) Kymographs show colocalization of kinesin and TRAK adaptors in the presence of 2 mM Ca^2+^. Assays were conducted in 10 nM MAP7. **g**, The percentage of KT colocalizers on MTs in the presence or absence of Ca^2+^ (*n* = 15, 15, 12, and 12 MTs from left to right, three independent trials). **h**, (Left) The Miro-binding model predicts that Ca^2+^ binding to Miro1 triggers the binding of the kinesin motor domain to Miro1 instead of MTs. (Right) Kymographs show that tail-truncated kinesin (KIF5B^1-490^) does not colocalize with Miro1 in the presence or absence of 2 mM Ca^2+^. Assays were performed in the absence of TRAK adaptors and MAP7. In (**c**), (**e**), and (**g**), the center line and whiskers represent mean and s.d., respectively, and p-values are calculated from a two-tailed t-test. Arrowheads show colocalization of Miro1 to processive kinesins in (**b**) and (**f**), and TRAK in (**d**).

We next tested whether longer TRAK1 or TRAK2 constructs could form a ternary complex with kinesin and Miro1. Co-immunoprecipitation assays showed that Miro1 interacts directly with either full-length TRAK1^1-953^ and TRAK2^1-914^, or a TRAK1 construct containing the coiled coils and part of the C-terminal domain (TRAK1^1-532^, **Extended Data Fig. 5f)**. These interactions are stable in the presence of a nonhydrolyzable GTP analog (GTPγS) or GDP, suggesting that Miro1’s nucleotide state does not affect its association with TRAK **(Extended Data Fig. 5f)**. We observed that TRAK1^1-532^ colocalizes with kinesin and recruits Miro1 to the complex in motility assays **(Fig. 5e,f)**, demonstrating that kinesin forms a stable complex with TRAK1 and Miro1.

### The mechanism of Ca^2+^-mediated arrest of the mitochondrial transport machinery

We used our in vitro reconstitution assay to test the predictions of the models that explain how Ca^2+^ binding to Miro1 might stall this transport machinery. Although the motor detachment model predicted that kinesin decouples from TRAK/Miro in excess Ca^2+^ **(Fig. 5f)**^16^, both KT and KTM complexes exhibited robust motility, and their landing rate and velocity were unaffected by the addition of 2 mM Ca^2+^ **(Fig. 5d and f, Supplementary Videos 6 and 7)**. We also tested the Miro-binding model by investigating whether Miro1 binds to the kinesin motor domain and prevents its motility in a Ca^2+^-dependent manner^18^ **(Fig. 5h)**. A kinesin construct that contains the motor domain but lacks the tail domain (KIF5B^1-490^) neither pulled down Miro1 in immunoprecipitation assays nor colocalized with Miro1 in the presence or absence of Ca^2+^ in motility assays **(Fig. 5h and Extended Data Fig. 6a-c)**. In the presence of Miro1, the run frequency of KIF5B^1-490^ remained unaffected by the addition of 2 mM Ca^2+^ **(Extended Data Fig. 6d)**. Collectively, our results indicate that mitochondrial pausing in response to elevated Ca^2+^ is not due to inhibition or disintegration of the KTM complex.

**Fig. 6.**
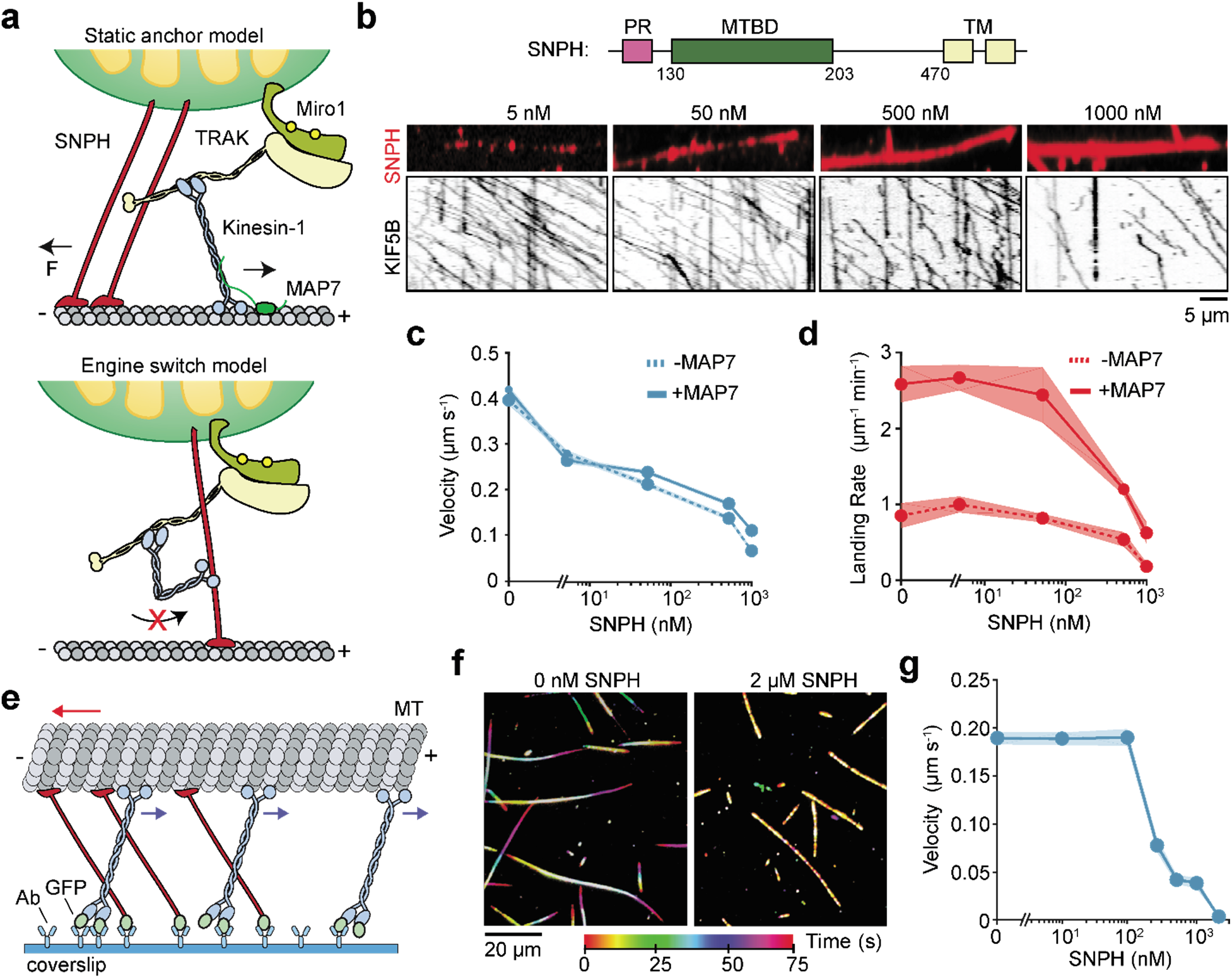
SNPH stalls MT gliding by multiple kinesin motors. **a**, Static anchor and engine switch models for SNPH-mediated stalling of mitochondrial transport. **b**, (Top) Domain organization of human SNPH (MTBD: MT-binding domain, PR: proline-rich domain). (Bottom) Sample kymographs of kinesin motility on SNPH-decorated MTs in the presence of 10 nM MAP7. **c**, The average velocity of kinesin in the presence (±s.e.m., *n* = 98, 158, 95, 58, and 20 from left to right) and absence of 10 nM MAP7 (*n* = 115, 191, 154, 93, and 59 from left to right). **d**, The landing rate of kinesin in the presence and absence of 10 nM MAP7 (mean ± s.e.m., *n* = 9 MTs for all time points in both conditions, three independent trials). **e**, Schematic of the MT gliding assay. Kinesins were fixed on the glass surface from their tail through a GFP-antibody linkage. MTs glide with their minus-ends in the lead (red arrow) due to the plus-end directed motility of kinesins (blue arrow). Static binding of SNPH to MTs exerts resistive forces against gliding motility. **f**, Representative color-coded time projections of Cy5-MTs in the presence or absence of 2 μM SNPH-sfGFP. **g**, Quantification of MT gliding velocities as a function of increasing SNPH-sfGFP concentration (mean ± s.e.m., *n* = 50, 57, 58, 59, 71, 70, 61 MTs from left to right, three independent trials).

Finally, we investigated whether the mitochondrial transport can be stalled through static anchoring to MTs or inactivating kinesin by SNPH **(Fig. 6a)**^22^. To test these models, we expressed an SNPH construct that lacks the C-terminal transmembrane domain (SNPH^1-473^, **Extended Data Fig. 7a**) and confirmed that it densely decorates the MT surface^21^ **(Fig. 6b)**. If SNPH binds and inhibits kinesin, decoration of the MT surface by SNPH would increase the kinesin landing rate, but prevent subsequent motility. However, SNPH substantially decreased the kinesin landing rate and slowed down motility **(Fig. 6b-d and Supplementary Video 8)**. We also did not observe strong interactions between kinesin and SNPH in immunoprecipitation assays **(Extended Data Fig. 7b).** These results indicate that SNPH does not inhibit kinesin motility through direct interactions, but its MT binding may serve as an obstacle that reduces MT recruitment and velocity of kinesin, similar to MT-associated proteins (MAPs)^44^.

To test whether SNPH can induce resistive forces against motility by statically anchoring mitochondria to MTs, we immobilized kinesin and SNPH to the glass surface and asked how MT binding of SNPH affects MT gliding activity of kinesin motors **(Fig. 6e)**. Consistent with SNPH serving as a static anchor to stall mitochondrial transport, the addition of SNPH slowed down gliding motility **(Fig. 6f-g and Supplementary Video 9)**. MT gliding was completely stopped at a ~100-fold molar excess of SNPH **(Fig. 6g and Extended Data Fig. 7c,d**), suggesting that multiple SNPH molecules may be required to efficiently counter the motility of kinesin motors.

## Discussion

In this study, we reconstituted the mitochondrial transport machinery and showed that Miro1 and TRAK1/2 recruit kinesin and dynein to form a minimal complex sufficient to drive transport in anterograde and retrograde directions. We used this assay to test the existing models on the regulation of motors on TRAK adaptors and Ca^2+^-mediated arrest of mitochondrial transport. Based on our results, we propose a comprehensive model for how mitochondria are transported from the cell body to distal regions by kinesin, remain stationary at regions with high neuronal activity, and recycled back to the cell body by dynein **(Fig. 7)**.

**Fig. 7.**
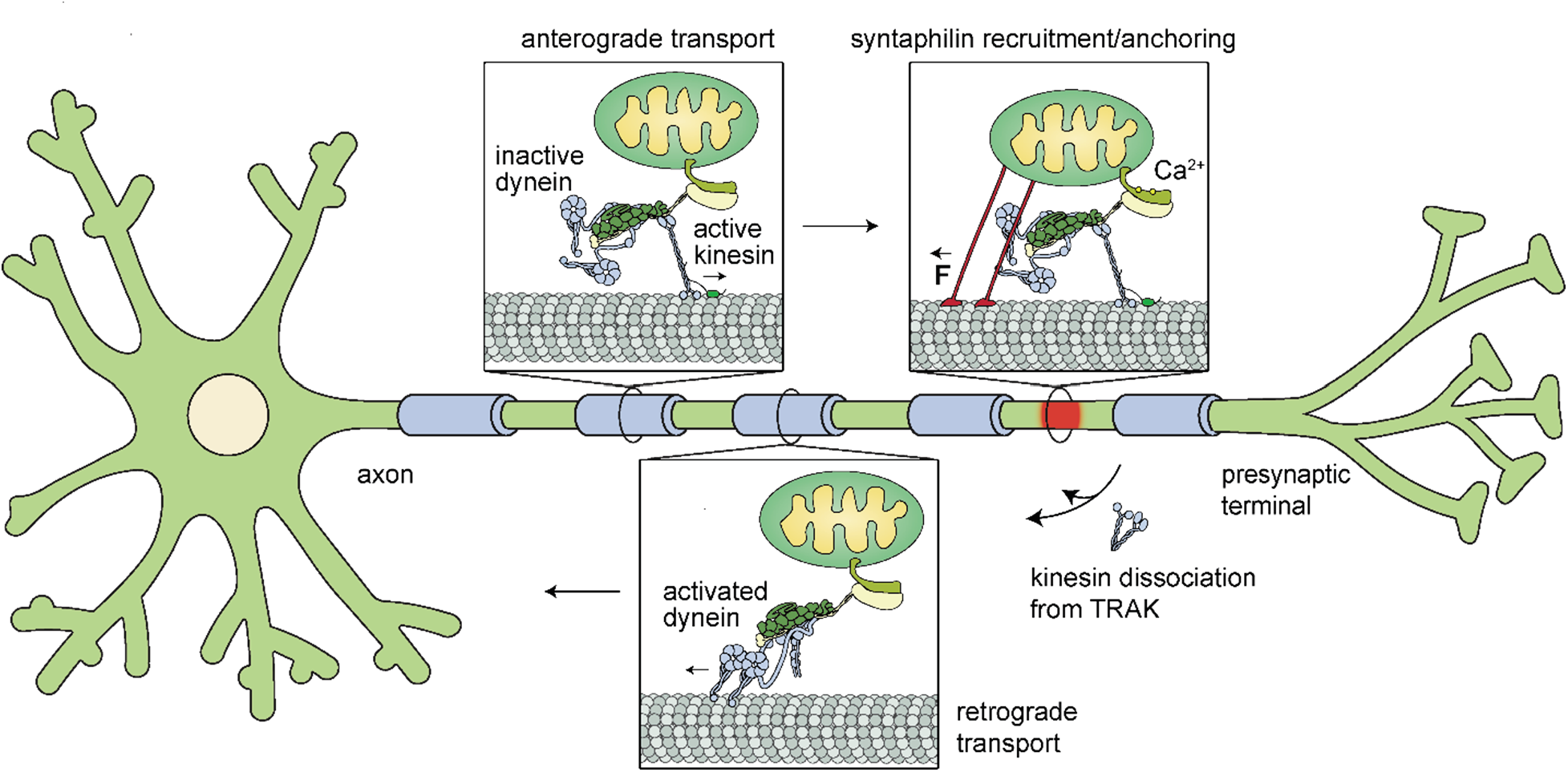
Model for bidirectional transport and Ca^2+^-mediated arrest of mitochondria in neurons. Mitochondria are transported anterogradely by active kinesin motors recruited by TRAK while dynein-dynactin is transported as an inactive passenger. In regions with high neuronal activity (red), mitochondria recruit SNPH, which anchors the mitochondria to the MT and stalls the transport machinery. Upon damage or stress, mitochondria are transported retrogradely for repair or clearance. Retrograde transport is initiated by dissociation of kinesin from TRAK, followed by activation of the dynein-dynactin-TRAK complex.

Specifically, TRAK1 and TRAK2 are bona fide adaptors that activate the motility and force generation of dynein-dynactin. Consistent with studies in neurons^14^, TRAK1 and to a lesser extent TRAK2, also recruits kinesin, but these interactions do not substantially activate the motor for processive motility. A recent in vitro reconstitution study also showed that Hook3 adaptors recruit kinesin-3 but do not activate its processive motility^41^. Therefore, cargo binding may not be sufficient for the full activation of kinesin motors. We observed that MAP7 decoration of MTs promoted the motility of KT complexes, highlighting the necessity of MAP7 for most, if not all, kinesin-1 driven transport in various cell types^38,45,46^.

Kinesin and dynein motors are interdependent on mitochondria because selective knockdown of kinesin disrupts transport in both retrograde and anterograde directions^3,47^. How mitochondrial adaptors control the direction of transport driven by these opposing motors has remained a major unsolved problem. Our results indicate that TRAK adaptors act as a scaffold that coordinates kinesin and dynein activity to control the direction of transport. TRAK adaptors that simultaneously recruit kinesin and dynein exclusively moved towards the plus-end at speeds comparable to TRAK driven only by kinesin. This is consistent with the observation that mitochondria move at similar speeds to kinesin in an anterograde direction^50^ even though dynein is localized to these cargos^48,49^. We propose that dynein is transported as an inactive passenger when kinesin drives anterograde transport of mitochondria **(Fig. 7)**. However, we did not observe dynein to transport kinesin towards the minus-end because the assembly of the active DDT complexes is mutually exclusive with kinesin binding **(Fig. 4e)**. These observations suggest that retrograde transport of mitochondria is initiated by dissociation of kinesin and formation of active dynein/dynactin on TRAK **(Fig. 7)**.

The molecular cues that govern the activation and inhibition of motors on TRAK adaptors remain to be investigated. We showed that both kinesin and dynein interact with the coiled-coil domain of TRAK. Because the coiled coils of adaptor proteins run along the length of the dynactin filament^51,52^, kinesin binding to TRAK may destabilize or partially overlap with the interface between dynactin and TRAK. As a result, kinesin binding may reduce the recruitment and activation of dynein/dynactin on TRAK adaptors. Interestingly, our results are markedly different from the motility of the reconstituted kinesin-3/Hook3/dynein-dynactin complex, in which dynein can transport kinesin-3 toward the minus-end^41^. Therefore, cargo-specific adaptor proteins may utilize distinct mechanisms to coordinate opposing motors.

Using our in vitro reconstitution assay, we provide critical insight into how elevated levels of intracellular Ca^2+^ arrest mitochondrial transport. the assembly and motility of the KTM complexes were unaffected by increased Ca^2+^ levels, which is incompatible with both ‘motor detachment’ and ‘Miro binding’ models^16,18^. In addition, Miro1 directly interacts with kinesin-1’s tail, and this interaction was disrupted in the presence of Ca^2+^ or TRAK adaptors. The physiological significance of this Miro-kinesin interaction remains unclear, but it could enable localization of kinesin to retrogradely moving mitochondria independent of TRAK in order to recycle these motors to the cell body.

Our results show that Ca^2+^-mediated docking of mitochondria occurs upstream of the motor transport machinery **(Fig. 7)**. In support of this, we provided evidence that the MT anchoring protein SNPH is sufficient to stall the MT gliding activity of kinesin motors. We propose that static anchoring of mitochondria to MTs by SNPH resists both plus- and minus-end directed motility, which provides an explanation for why Ca^2+^-mediated recruitment of SNPH arrests both anterogradely and retrogradely moving mitochondria in neurons.

Collectively, our work provides a molecular explanation for how opposing actions of kinesin and dynein are coordinated by TRAK to control the directionality of mitochondrial transport and how the motor machinery is arrested at regions of elevated Ca^2+^. Mitochondrial transport is regulated by a plethora of other factors in cells. In particular, the dynein-interacting protein Disrupted in Schizophrenia 1 (DISC1) ^53^, Armadillo repeat-containing X-linked (Armcx) 1 and 3^54,55^, and mitochondrial fusion proteins MFN1 and MFN2 have been shown to interact with the Miro1/TRAK complex, and the genetic knockdown of these factors led to defects in mitochondrial transport ^1^. The in vitro reconstitution assay we developed in this study provides an experimental platform to investigate the molecular basis of how these factors regulate mitochondrial transport in cells.

## Methods

### Cloning and plasmid generation

The construct expressing the phi-dynein mutant (SNAP–DHC E1518K/R1567K), full-length TRAK1, and full-length TRAK2 were provided in a pACEBac1 vector backbone by A.P. Carter (MRC, University of Cambridge). The sequences encoding fulllength or truncated versions of human TRAK1 and TRAK2 were cloned into the pOmniBac vector. All constructs contained an N-terminal His6-ZZ tag followed by a TEV protease cleavage site for protein purification and a C-terminal SNAP-tag or GFP fusion for labeling and imaging purposes. A cDNA for full-length human kinesin (*KIF5B*; amino acids 1–963, clone ID 8991995) was obtained from GE Dharmacon and fused to GFP-SNAPf at its C terminus. The phi mutant of the dynein-1 heavy chain (DHC; SNAP-DHC E1518K/R1567K) was co-expressed by fusing the coding sequence to the pDyn2 plasmid containing genes encoding IC2C, LIC2, TCTEX1, LC8, and ROBL1, as described^30^. The list of constructs used for each dataset is given in **Supplementary Table 1**.

### Protein expression and purification

Native dynactin was purified from pig brains with the large-scale SP-Sepharose protocol and anion-exchange chromatography using a MonoQ column (GE), as described previously^56^.

The phi mutant of dynein, KIF5B, MAP7, TRAK1, and TRAK2 constructs were purified from baculovirus-infected SF9 cells. Briefly, cell pellets were resuspended in a lysis buffer (see below) supplemented with 1x protease inhibitor cocktail (Roche cOmplete Protease Inhibitor Cocktail) and lysed using a Dounce homogenizer (20 strokes with loose plunger followed by 20 strokes tight plunger). The cell lysate was clarified by centrifugation at 186,000 g for 45 min, incubated with IgG Sepharose (GE Healthcare) for 2 hr at 4°C, applied to a gravity flow column, and washed extensively with a TEV wash buffer (see below). The protein-bead complexes were then treated with TEV protease at 12°C overnight. The mixture was then centrifuged at 4000 rpm for 5 min and the supernatant was concentrated using an Amicon Ultra-0.5 mL spin column (EMD Millipore). Protein concentration was determined by measuring the OD_280_ using Nanodrop 1000.

Different buffer conditions were used for each protein preparation. Cells expressing dynein were lysed in a dynein lysis buffer (50 mM HEPES pH 7.4, 100 mM NaCl, 10% Glycerol, 1mM DTT, 2 mM PMSF), IgG beads were washed with a dynein TEV wash buffer (50 mM Tris-HCl pH 7.4, 150 mM K-Acetate, 2 mM Mg-Acetate, 1 mM EGTA, 10% Glycerol, 1mM DTT) and the protein was concentrated using a 100K molecular weight cut-off (MWCO) spin filter. For kinesin purification, the cells were lysed in a kinesin lysis buffer (50 mM HEPES pH 7.4, 1 M NaCl, 10% Glycerol, 1mM DTT, 2 mM PMSF), IgG beads were washed with a kinesin TEV wash buffer (50 mM HEPES pH 7.4, 300 mM NaCl, 10% Glycerol, 1mM DTT), and the protein was concentrated using 100K MWCO spin filter. MAP7 purification was performed using a MAP7 lysis buffer (25 mM HEPES pH 7.4, 1 M KCl, 10% Glycerol, 1mM DTT, 1 mM PMSF) and MAP7 TEV wash buffer (25 mM HEPES pH 7.4, 300 mM KCl, 1 mM EGTA, 10 mM MgCl_2_, 10% Glycerol, 1mM DTT). MAP7 was concentrated using a 50K MWCO spin filter.

Truncated TRAK constructs were purified in the presence of high salt, glutamic acid, and arginine to improve solubility (TRAK Lysis buffer: 50 mM HEPES pH 7.4, 1M NaCl, 10% Glycerol, 50 mM L-Glu, 50 mM L-Arg, 1mM DTT, 2 mM PMSF; and TRAK TEV wash buffer: 50 mM HEPES pH 7.4, 500 mM NaCl, 50 mM L-Glu, 50 mM L-Arg, 10% Glycerol, 1mM DTT), and protein was concentrated using a 50K MWCO spin column. The concentrated protein solution was then dialyzed overnight at 4 °C into the wash buffer to remove amino acids in 50 mM HEPES pH 7.4, 300 mM NaCl, 20% glycerol, 1 mM DTT. Full-length TRAK1 was purified from baculovirus-infected SF9 cells while full-length TRAK2 was purified from baculovirus-infected HEK 293F GNTI^-/-^ cells using the same purification method, except the lysis buffer was additionally supplemented with 0.1% Triton X-100.

Miro1^1-592^ (Miro1^1-592^-SNAP-psc-StrepII) was purified from baculovirus-infected SF9 cells. Cells were resuspended in Miro lysis buffer (25 mM HEPES pH 7.4, 300 mM NaCl, 10% Glycerol, 1mM DTT, 1 mM PMSF) supplemented with 1x protease inhibitor cocktail and lysed using a Dounce homogenizer. The lysate was clarified by centrifugation (65,000 rpm for 45 min) and incubated with Streptactin Sepharose (IBA) for 2 hr at 4°C, applied to a gravity flow column, and washed extensively with Miro wash buffer (25 mM HEPES pH 7.4, 300 mM NaCl, 10% Glycerol, 1mM DTT). The protein was then eluted from beads with 3 mM desthiobiotin and concentrated using a 50K MWCO spin column.

SNPH^1-473^ was purified from BL21(DE3) *E. coli* cells. Briefly, cell pellets were resuspended in SNPH lysis buffer (25 mM HEPES pH 7.4, 300 mM NaCl, 10% Glycerol, 1mM DTT, 1 mM PMSF) supplemented with 1x protease inhibitor cocktail and lysed using a sonicator for 2 min. The lysate was clarified by centrifugation (65,000 rpm for 45 min) and incubated with IgG Sepharose for 2 hr at 4°C, applied to a gravity flow column, and washed extensively with wash buffer (25 mM HEPES pH 7.4, 300 mM NaCl, 10% Glycerol, 1mM DTT). The protein-bead complexes were then treated with TEV protease at 4°C overnight. The mixture was then centrifuged at 4000 rpm for 5 min and the supernatant was concentrated using a 50K MWCO spin column.

### Labeling

Proteins were labeled with fluorescent probes before they were eluted from the affinity columns. For SNAP labeling, bead slurry was concentrated to 5 mL, followed by the addition of 5 nmol of either BG-LD555 or BG-LD655 dye, followed by incubation for 1 h at 4 °C. The slurry was added to a gravity flow column and washed extensively in a wash buffer. For ybbR labeling, bead slurry was concentrated to 5 mL, followed by the addition of 5 nmol of either CoA-LD555 or BG-LD655 dye followed by incubation for 30 min at room temperature in the presence of 1 μM Sfp phosphopantetheinyl transferase to catalyze protein labeling.

### Motility assays

Biotinylated MTs were prepared by mixing 98% unlabeled and 2% biotinylated pig brain tubulin in BRB80 supplemented with 2 mM GTP and 20% DMSO for 30 min at 37 °C. MT polymerization was stabilized by 100 nM taxol and the reaction was incubated for an additional 60 min. Unpolymerized tubulin was removed by pelleting at 20,000*g* for 12 min and resuspending MTs in BRB80 containing 100 nM taxol.

To immobilize biotinylated MTs to the coverslip, 1 mg ml^−1^ BSA-biotin (Sigma) was introduced into the flow chamber, which was then washed with 1x MB buffer (30 mM HEPES, 5 mM MgSO4, 1 mM EGTA, pH 7.0) supplemented with 1 mM DTT, 10 μM taxol, 1.25 mg ml^−1^ casein (Sigma) and 0.5% pluronic (MBCT). The chamber was incubated with 1 mg ml^−1^ streptavidin (NEB) and washed with MBCT. For imaging dynein motility, fluorescently-labeled dynein, dynactin, and a cargo adaptor (TRAK1 or TRAK2) were mixed at a 1:5:20 molar ratio in DLB. For imaging kinesin motility, fluorescently-labeled kinesin, a cargo adaptor (TRAK1 or TRAK2), MAP7, and Miro1^1-592^ were mixed at a 1:5:5:10 molar ratio in MB buffer. For imaging both dynein and kinesin simultaneously, fluorescently-labeled dynein, dynactin, kinesin, a cargo adaptor (TRAK1 or TRAK2), and MAP7 were mixed at a 1:5:1:1:5 ratio in MB buffer. The mixture was incubated on ice for 10 min and diluted 30-fold in MBC. Finally, the mixture was diluted 10-fold in the stepping buffer (MBCT supplemented with 0.1 mg ml^−1^ glucose oxidase, 0.02 mg ml^−1^ catalase, 0.8% D-glucose, and 1 mM Mg · ATP) and introduced into the chamber. Motility was recorded for 5 min. For assays including Miro1^1-592^, 0.1 mg ml^−1^ biotin-BSA was also included in the chamber for surface passivation.

### Gliding assays

For MT gliding assays, rabbit monoclonal anti-GFP antibody (~0.4 mg ml^−1^, Covance) was flown into an assay chamber and incubated for 3 min. The chamber was washed with 30 μl of MB supplemented with 1mM DTT, 10 μM taxol, and 1.25 mg ml^−1^ casein (Sigma). Subsequently, 10 μl of 2.5 nM GFP-tagged kinesin was added to the chamber. After 2 min incubation, the unbound motor was removed by washing the chamber with 30 μl MB. For experiments with SNPH, 10 μl of SNPH^1-473^-sfGFP was added to the chamber at the indicated concentration for 2 min followed by a 30 μl MB wash. Then, 10 μl of 200 nM Cy5-labeled MTs were flown to the chamber and allowed to bind the kinesin-decorated surface for 4 min. The chamber was then washed with 60 μl MB. Lastly, 10 μl of imaging buffer (MB supplemented with 0.02 mg ml^−1^ catalase, 0.8% D-glucose, and 1 mM Mg · ATP) was flown into the chamber to initiate gliding motility.

### Microscopy

Fluorescence imaging experiments were performed using a Nikon multicolor TIRF microscope equipped with a Nikon Ti-E microscope body, a 100X magnification 1.49 N.A. apochromat oil-immersion objective (Nikon), a perfect focusing system, and an electron-multiplied charge-coupled device camera (Andor, Ixon EM+, 512 × 512 pixels) with an effective pixel size of 160 nm after magnification. Alexa488/GFP, LD555, and LD655 probes were excited with 0.05 kW cm^−2^ 488-nm, 561-nm, and 633-nm laser beams (Coherent), and their fluorescent emissions were filtered through a notch dichroic filter and 525/40, 585/40, and 655/40 bandpass emission filters (Semrock), respectively. Multicolor fluorescence imaging was performed using the time-sharing mode in MicroManager. Videos were recorded at 2-4 Hz.

### Optical trapping assays

DDT_1_^1-400^ and DDT_2_^1-400^ complexes were assembled with 1 μL of 0.84 mg mL^−1^ dynein, 1 μL of 1.7 mg mL^−1^ dynactin, and 1 μL of 0.22 mg mL^−1^ of TRAK1^1-400^ or 0.1 mg mL^−1^ of TRAK2^1-400^ in DLB for 5 min at 4 °C. The protein mixture was then added to 700 nm diameter polystyrene beads coated with a polyclonal GFP antibody (Covance) and incubated for 10 minutes. Similarly, kinesin-TRAK1^1-400^ complexes were assembled with 1 μL of 0.05 mg · mL^−1^ kinesin and 0.1 mg · mL^−1^ of TRAK1^1-400^ before being added to the beads. Flow chambers were first decorated with Cy5-labeled sea urchin axonemes in 1x MB buffer. The motor-bead mixture was introduced to the chamber in the imaging buffer. To ensure that more than ~95% of beads were driven by single motors, the protein mixture was diluted before incubating with beads such that a maximum of 30% of beads exhibited activity when brought into contact with an axoneme.

Optical trapping experiments were performed on a custom-built optical trap microscope set-up^57^. Briefly, motor-coated beads were trapped with a 2 W 1,064-nm laser beam (Coherent) focused on the image plane using a 100X magnification 1.49 N.A. apochromat oil-immersion objective (Nikon). Cy5-labeled sea urchin axonemes were excited with a 633-nm HeNe laser (JDSU Uniphase), imaged using a monochrome camera (The Imaging Source), and moved to the center of the field of view using a locking XY stage (M-687, Physik Instrumente). The trapped bead was lowered to the surface of the axonemes using a piezo flexure objective scanner (P-721 PIFOC, Physik Instrumente). Bead position relative to the center of the trap was monitored by imaging the back-focal plane of a 1.4 N.A. oil-immersion condenser (Nikon) on a position-sensitive detector (First Sensor). Beam steering was controlled with a pair of perpendicular acousto-optical deflectors (AA Opto-Electronic). For calibrating the detector response, a trapped bead was rapidly raster-scanned by the acousto-optical deflector and trap stiffness was derived from the Lorentzian fit to the power spectrum of the trapped bead. The laser power was adjusted with a half-wave plate on a motorized rotary mount. The spring constant was set to ~0.04 pN nm^−1^ for DDT_1_^1-400^ and DDT_2_^1-400^, and ~0.08 pN nm^−1^ for kinesin-TRAK1 experiments.

Custom MATLAB software was used to extract stall forces and stall times from raw traces. First, raw traces were down-sampled from 5,000 Hz to 250 Hz. Stall events were defined as a stationary period of a motor at forces above 2.5 pN lasting a minimum of 100 ms, followed by snapping back of the bead to the trap center. The stall force was defined as the mean force in the last 20% for the stall event. The stall time was defined as the interval the bead spent at a force of at least 80% of the stall force. All stall events were plotted and manually reviewed to confirm the accuracy of the reported values.

### Data Analysis

Videos were analyzed in ImageJ. Kymographs were generated by plotting segmented lines along the MTs using a custom-written ImageJ macro. The processive movement was defined and analyzed as described previously^32^. Complexes that exhibited diffusive movement, ran for less than 250 nm, and paused for more than 1 s were excluded from velocity analysis. For two-color imaging, the fluorescence channels were overlaid in ImageJ to generate a composite image. Colocalization events were manually scored in kymographs. For two-color imaging of a motor and a cargo adaptor (TRAK1 or TRAK2), processive motility events observed in the cargo adaptor channel that did not colocalize with a motor were still included in the velocity analysis.

### Statistics and reproducibility

At least three independent repetitions were performed to obtain any given result. The number of replicates (*n*) and statistical analysis methods are clearly stated in the main text or the figure legends. Representative data are shown from independently repeated experiments.

## Data availability

All data that support the conclusions are available from the authors on request.

## Author contributions

J.C. and A.Y. conceived the study and designed the experiments. J.C. prepared the constructs and isolated the proteins. J.C. labeled the proteins with fluorescent dyes and performed the motility experiments. A.H. performed optical-trapping assays. J.C. and A.Y. wrote the manuscript, and all authors read and commented on the manuscript.

## Acknowledgments

We are grateful to the members of the Yildiz laboratory for helpful discussions and carefully reading the manuscript, Andrew Carter and Sami Chaaban (MRC, Cambridge) for providing plasmids and dynactin protein, Simon Bullock (MRC, Cambridge) for providing Lis1 protein, and Scott Blanchard (St. Jude’s Hospital) for synthesizing fluorescent dyes. This work was funded by grants from the NIH (GM094522), and NSF (MCB-1055017, MCB-1617028) to A.Y.

## Extended Data Tables

**Extended Data Table 1.** The list of constructs used in this study. The SNAPf-Dyn-Phi construct was a generous gift from A. Carter (MRC, UK). Other constructs were cloned into the pOmniBac vector for baculovirus expression in SF9 cells, except SNPH constructs which were cloned into the pET17b vector for bacterial expression. A ZZ-Tev tag was inserted at the N-termini of the constructs for binding the protein to IgG beads. The ZZ tag was cleaved by Tev protease to elute the protein from the beads. The SNAPf tag was used to label the proteins with fluorescent dyes or biotin functionalized with benzyl guanine. The ybbR tag was for SFP-catalyzed labeling of proteins with fluorescent dyes or biotin functionalized with CoA. A StrepII tag was inserted at the C-termini of Miro1^1-592^ and the construct was eluted with desthiobiotin (ED: Extended Data).

| Construct Name | Description | Figures |
| --- | --- | --- |
| SNAPf-Dyn-Phi | ZZ-Tev-SNAPf-DYNH1C1<sub>K1610/R1567E</sub>-IC2C-LIC2-Robl1-Tctex1-LC8 | Fig. 1c-f; Fig. 3; Fig. 4a-f; ED Fig. 1b-e; 3b-d; 4a,b |
| TRAK1<sup>1-360</sup>-SNAPf | ZZ-Tev-TRAK1<sup>1-400</sup>-SNAPf | Fig. 1c,e; Fig. 2a-c; Fig. 5d-g; ED Fig. 1a,b; 2b,c; 5d |
| TRAK1<sup>1-400</sup>-SNAPf | ZZ-Tev-TRAK1<sup>1-400</sup>-SNAPf | Fig. 1c,e,f; Fig. 3; Fig. 4a-c; ED Fig. 1a-d; Fig. 2d; 4a,b |
| TRAK2<sup>1-360</sup>-SNAPf | ZZ-Tev-TRAK2<sup>1-360</sup>-SNAPf | Fig. 1d,e; 5e,g; ED Fig. 1a,b, 2b,c; 5c,d, |
| TRAK2<sup>1-400</sup>-SNAPf | ZZ-Tev-TRAK2<sup>1-400</sup>-SNAPf | Fig. 1d-f; 3; Fig. 4a,b; ED Fig. 1a-c,e; Fig. 2d; 4a,b |
| KIF5B-GFP-SNAPf | ZZ-Tev-KIF5B-GFP-SNAPf | Fig. 2a-c; 3B; Fig. 4a,b; Fig. 5b-g; Fig. 6f,g; ED Fig. 2a-f; 4a,b; 5b-d; 7b |
| MAP7-ybbR | ZZ-Tev-MAP7-ybbR | Fig. 2a-c; Fig. 4a-b; Fig. 5b-g; Fig. 6b-d; ED Fig. 2a,c,d-f; 4a,b; 5c,d; 7b |
| TRAK1<sup>1-360</sup>-GFP | ZZ-Tev-TRAK1<sup>1-360</sup>-GFP | ED Fig. 3a,b |
| TRAK1<sup>1-400</sup>-GFP | ZZ-Tev-TRAK1<sup>1-400</sup>-GFP | Fig. 3; ED Fig. 3a-d |
| TRAK2<sup>1-360</sup>-GFP | ZZ-Tev-TRAK2<sup>1-360</sup>-GFP | ED Fig. 3a,b |
| TRAK2<sup>1-400</sup>-GFP | ZZ-Tev-TRAK2<sup>1-400</sup>-GFP | Fig. 3; ED Fig. 3a-d |
| TRAK1<sup>1-532</sup>-SNAPf | ZZ-Tev-TRAK1<sup>1-532</sup>-SNAPf | Fig. 5d,e; ED Fig. 5a,e |
| TRAK2<sup>1-532</sup>-SNAPf | ZZ-Tev-TRAK2<sup>1-532</sup>-SNAPf | Fig. 5e; ED Fig. 5a,c |
| KIF5B-ybbR | ZZ-Tev-KIF5B-ybbR | Fig. 3; Fig. 6a; ED Fig. 3a,c,d |
| KIF5B<sup>Δ1-336</sup>-GFP-SNAPf | ZZ-Tev-KIF5B<sup>Δ1-336</sup>-GFP-SNAPf | Fig. 4c,d; ED Fig. 2a |
| KIF5B<sup>1-490</sup>-GFP-SNAPf | ZZ-Tev-KIF5B<sup>1-490</sup>-GFP-SNAPf | Fig. 5h; ED Fig. 6a-d |
| Miro1<sup>1-592</sup>-SNAPf-psc-StrepII | ZZ-Tev-Miro1<sup>1-592</sup>-SNAPf-psc-StrepII | Fig. 5b-h; ED Fig. 5a-e; 6b-d |
| TRAK1<sup>1-953</sup>-sfGFP-SNAPf | ZZ-Tev-TRAK1<sup>1-953</sup>-sfGFP-SNAPf | ED Fig. 2a,e,f; 5e |
| TRAK2<sup>1-914</sup>-SNAPf | ZZ-Tev-TRAK2<sup>1-914</sup>-SNAPf | ED Fig. 2a,e,f; 5e |
| SNPH<sup>1-473</sup>-sfGFP | ZZ-Tev- SNPH<sup>1-473</sup>-sfGFP | Fig. 6c-g; ED Fig. 7a,c,d |
| SNPH<sup>1-473</sup>-ybbR | ZZ-Tev- SNPH<sup>1-473</sup>-ybbR | ED Fig. 7a,b |

## Extended Data Figures

**Extended Data Fig. 1.**
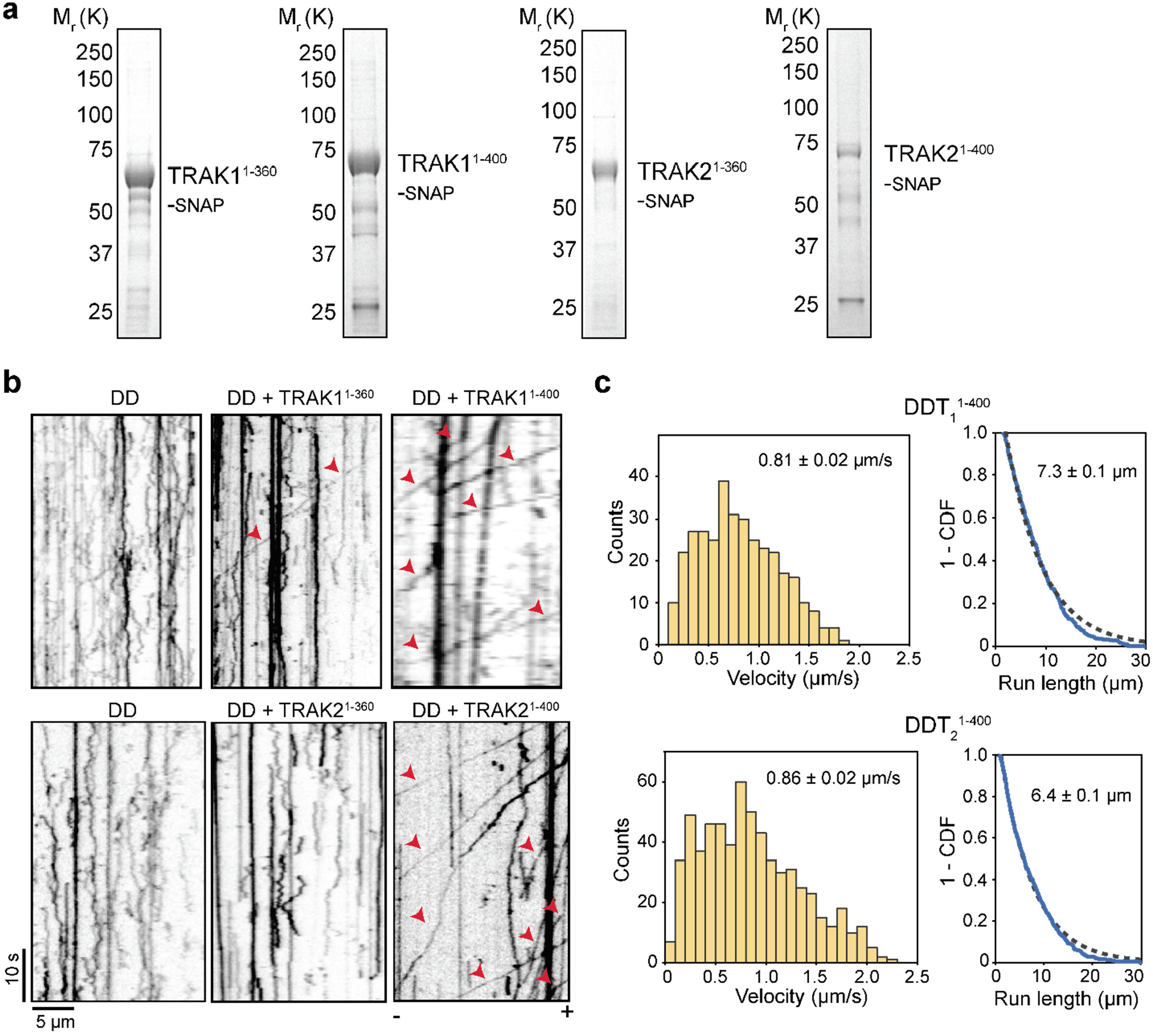
Purification and motility of DDT complexes. **a**, Denaturing gel pictures of purified TRAK constructs. **b**, Sample kymographs of LD655-dynein in the presence of dynactin and TRAK1 or TRAK2 constructs in 2 mM ATP. Arrowheads show mobile complexes. **c**, Velocity histograms (mean ± s.e.m.) and the inverse cumulative distributions (1-CDF) of motor run length for DDT_1_^1-400^ (*n* = 341 molecules from three independent experiments) and DDT_2_^1-400^ (*n* = 417 molecules from three independent experiments). Fits to a single exponential decay (dashed curves) reveal the mean run lengths (±s.e.).

**Extended Data Fig. 2.**
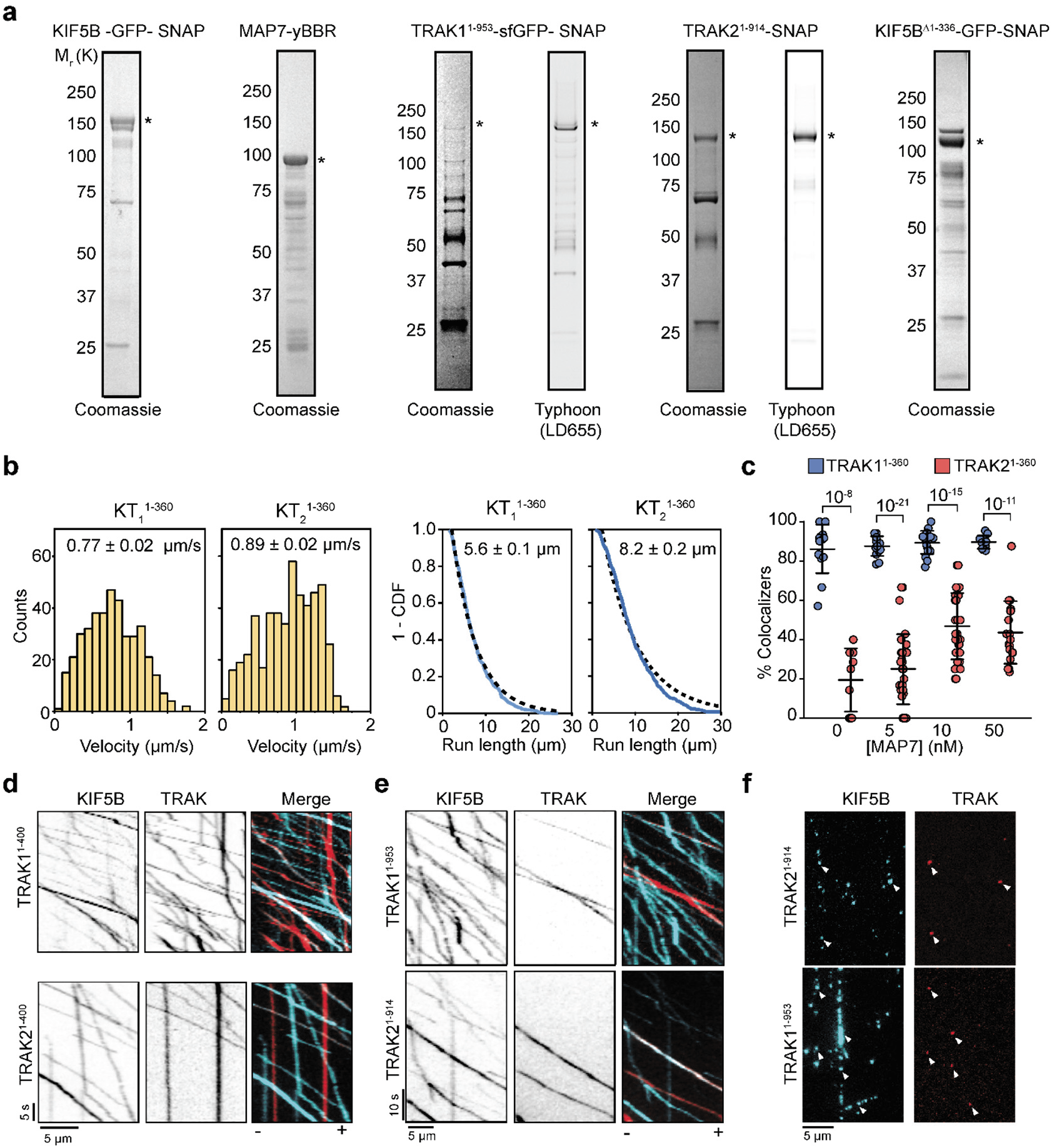
Purification and motility of KT complexes. **a**, Denaturing gel pictures of purified kinesin (KIF5B), MAP7, and TRAK constructs. Stars indicate the expected molecular weight. **b**, (Left) Velocity histogram (mean ± s.e.m.) and (Right) 1-CDF of motor run length for KT_1_^1-360^ (*n* = 404) and KT_2_^1-360^ (*n* = 236, three independent experiments). Fits to a single exponential decay (dashed curves) reveal the mean run lengths (±s.e.). **c**, The percentage of TRAK colocalization to processive kinesin motors moving along the MTs in the presence or absence of MAP7. The center line and whiskers represent mean and s.d., respectively (*n* = 13, 14, 18, 12, 9, 22, 29, and 19 MTs from left to right, three independent trials). P values are calculated from a two-tailed t-test. **d**, Sample kymographs of LD555-kinesin and LD655-labeled TRAK1^1-400^ or TRAK2^1-400^ in 10 nM MAP7. **e**, Sample kymographs of LD555-kinesin and LD655-labeled full-length TRAK1 ^1-953^ or TRAK2^1-914^ in 10 nM MAP7. **f**, Sample frames of LD555-kinesin (cyan) LD655-labeled TRAK1^1-953^ or TRAK2^1-914^ (red) in the presence of 10 nM MAP7. White arrows indicate colocalizers of both kinesin and TRAK.

**Extended Data Fig. 3.**
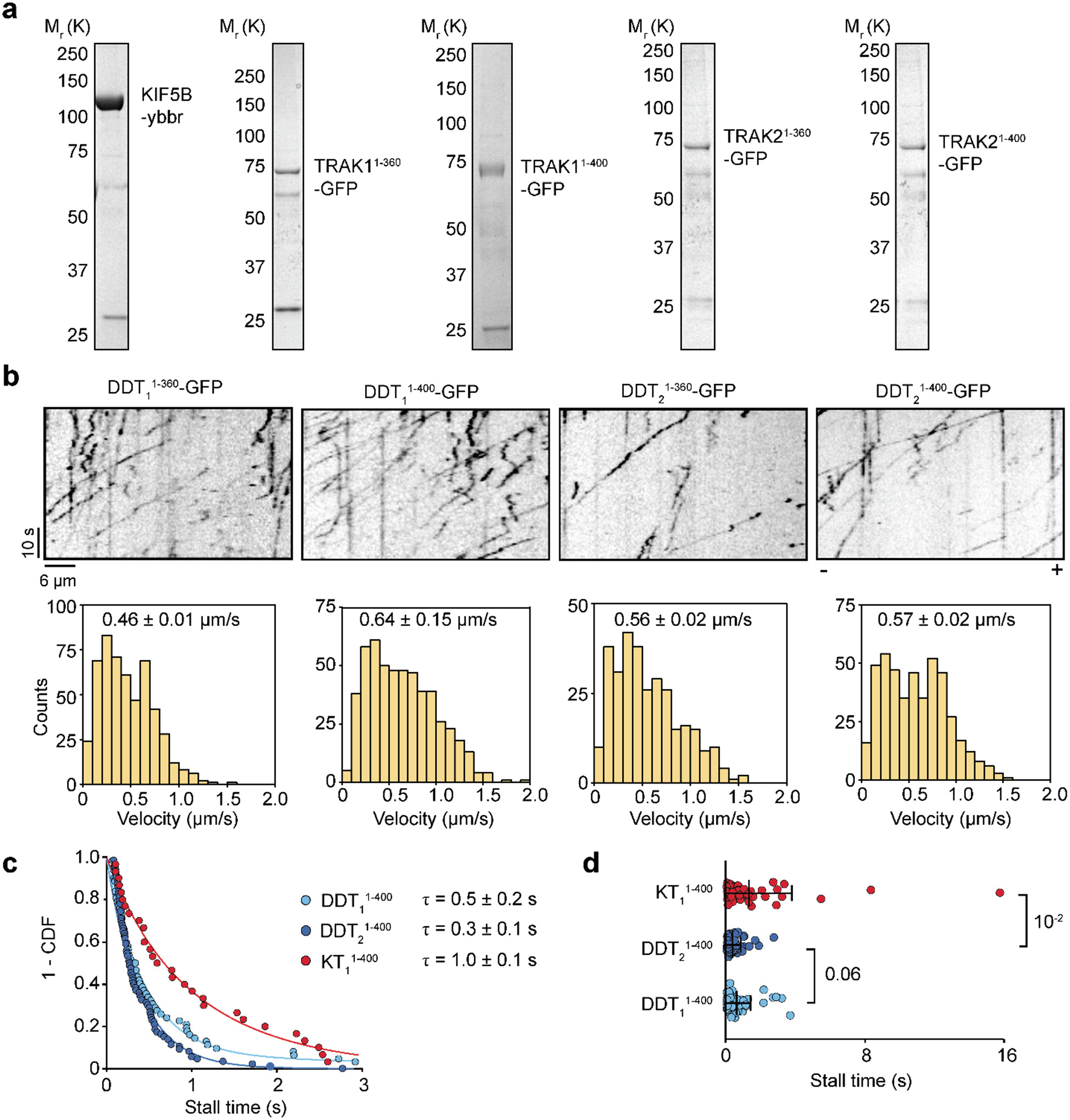
Purification and motility of DDT complexes for optical trapping experiments. **a**, Denaturing gel pictures of purified KIF5B-ybbR, and GFP-tagged TRAK constructs. **b**, (Top) Sample kymographs representing the motility of DDT complexes in 2 mM ATP. (Bottom) Velocity histograms of DDT complexes in 2 mM ATP (mean ± s.e.m., *n* = 524, 523, 312, and 454 from left to right, three independent experiments per condition). **c**, 1-CDF of motor stall times. Mean stall times (τ, ± s.e.) were calculated from a fit to a single exponential decay (solid curves). **d**, Box plots of stall times for DDT_1_^1-400^, DDT_2_^1-400^, and KT_1_^1-400^ complexes Center line and whiskers show the mean and s. d. P-values are calculated from a two-tailed t-test. In (**c**) and (**d**), *n* = 62, 86, and 55 for KT_1_^1-400^, DDT_2_^1-400^, and DDT_1_^1-400^, respectively.

**Extended Data Fig. 4.**
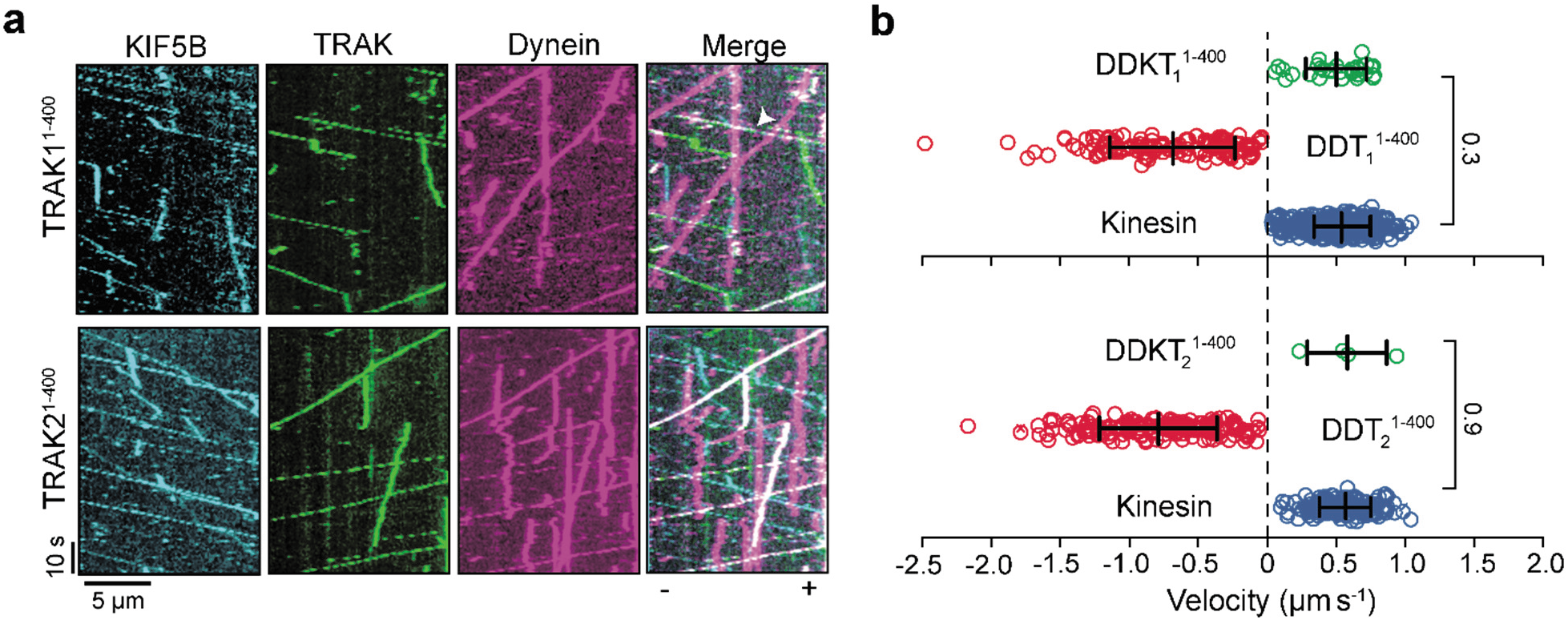
Motility of DDKT complexes in the presence of Lis1. **a**, Sample kymographs of Alexa488-kinesin, LD555-TRAK, and LD655-dynein on a surface-immobilized microtubule in the presence of unlabeled dynactin, 5 nM MAP7 and 1 μM Lis1. White arrowhead shows colocalization of kinesin, dynein, and TRAK. **b**, The velocity distribution of individual motor complexes and DDKT assemblies in the presence of 1 μM Lis1. Negative velocities correspond to minus-end-directed motility (*n* = 30, 140, 439, 4, 175, and 165 from top to bottom, with three independent experiments per condition). The center line and whiskers represent the mean and s.d., respectively. P-values are calculated from a two-tailed t-test.

**Extended Data Fig. 5.**
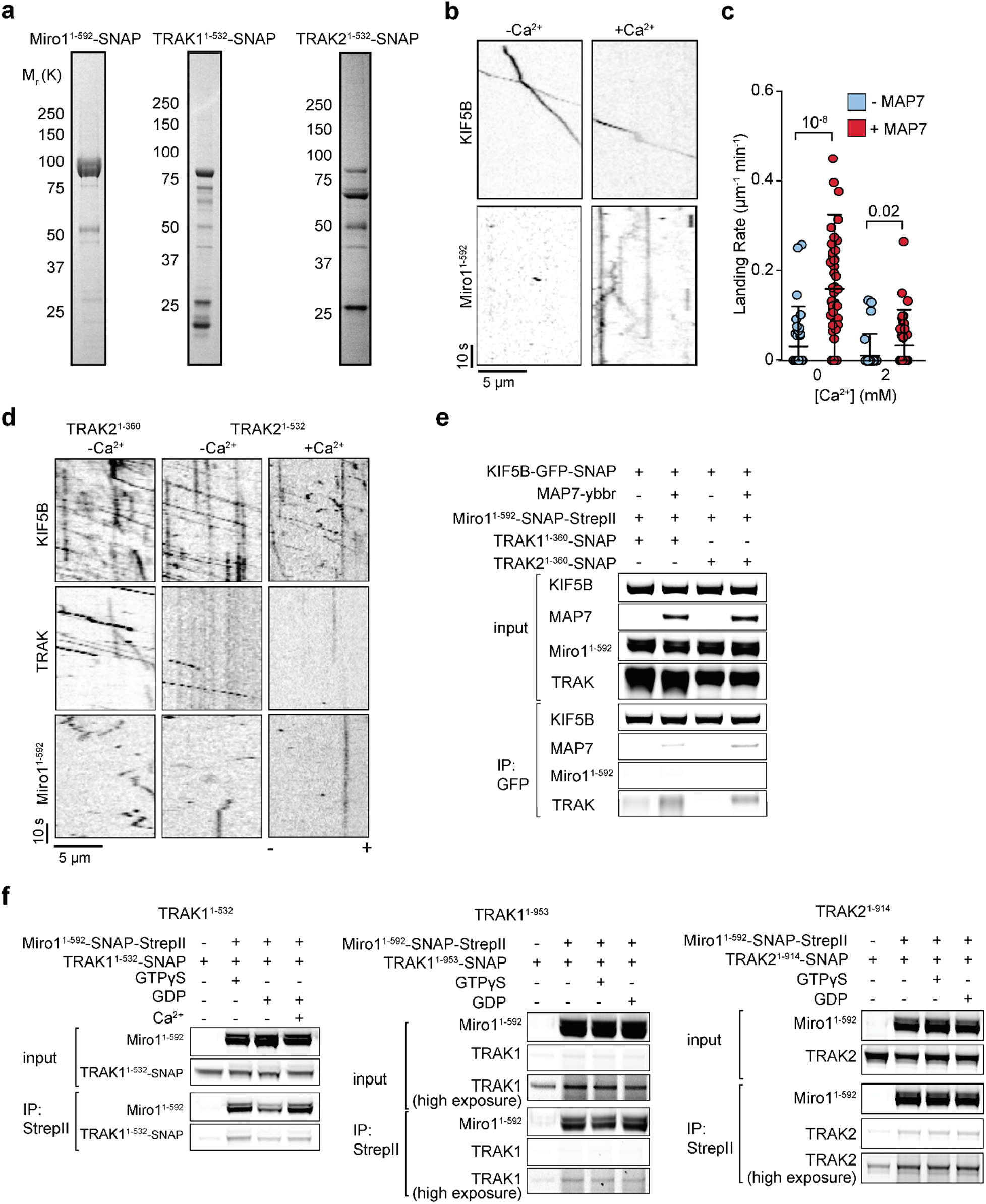
Miro1 interacts with TRAK under different nucleotide conditions both in the presence and absence of Ca^2+^. **a**, Denaturing gel pictures of purified Miro1^1-592^-SNAP, TRAK1^1-532^-SNAP, and TRAK2^1-532^-SNAP. **b**, Kymographs of LD655-kinesin (KIF5B) and LD555-Miro1^1-592^ in the absence and presence of 2 mM Ca^2+^. Assays were conducted without MAP7. **c**, The landing rate of kinesin-Miro1 co-localizers in the presence or absence of 10 nM MAP7 at 0 and 2 mM Ca^2+^ (*n* = 47, 43, 45, and 50 MTs from left to right, three independent trials). **d**, Kymographs of Alexa488-KIF5B, LD655-Miro1^1-592^, and LD555-labeled TRAK2 constructs. Assays were conducted in the presence of 10 nM MAP7. **e**, In vitro immunoprecipitation of purified KIF5B, TRAK1^1-360^ or TRAK2^1-360^, Miro1^1-592^, and MAP7. **f**, In vitro immunoprecipitation of purified, Miro1^1-592^ and either TRAK1^1-532^ (left), full-length TRAK1 (middle), or full-length TRAK2 (right). KIF5B-GFP-SNAP was present in all conditions. Assays were conducted in the presence of 100 μM GTPγS, 100 μM GDP, or 2 mM Ca^2+^.

**Extended Data Fig. 6.**
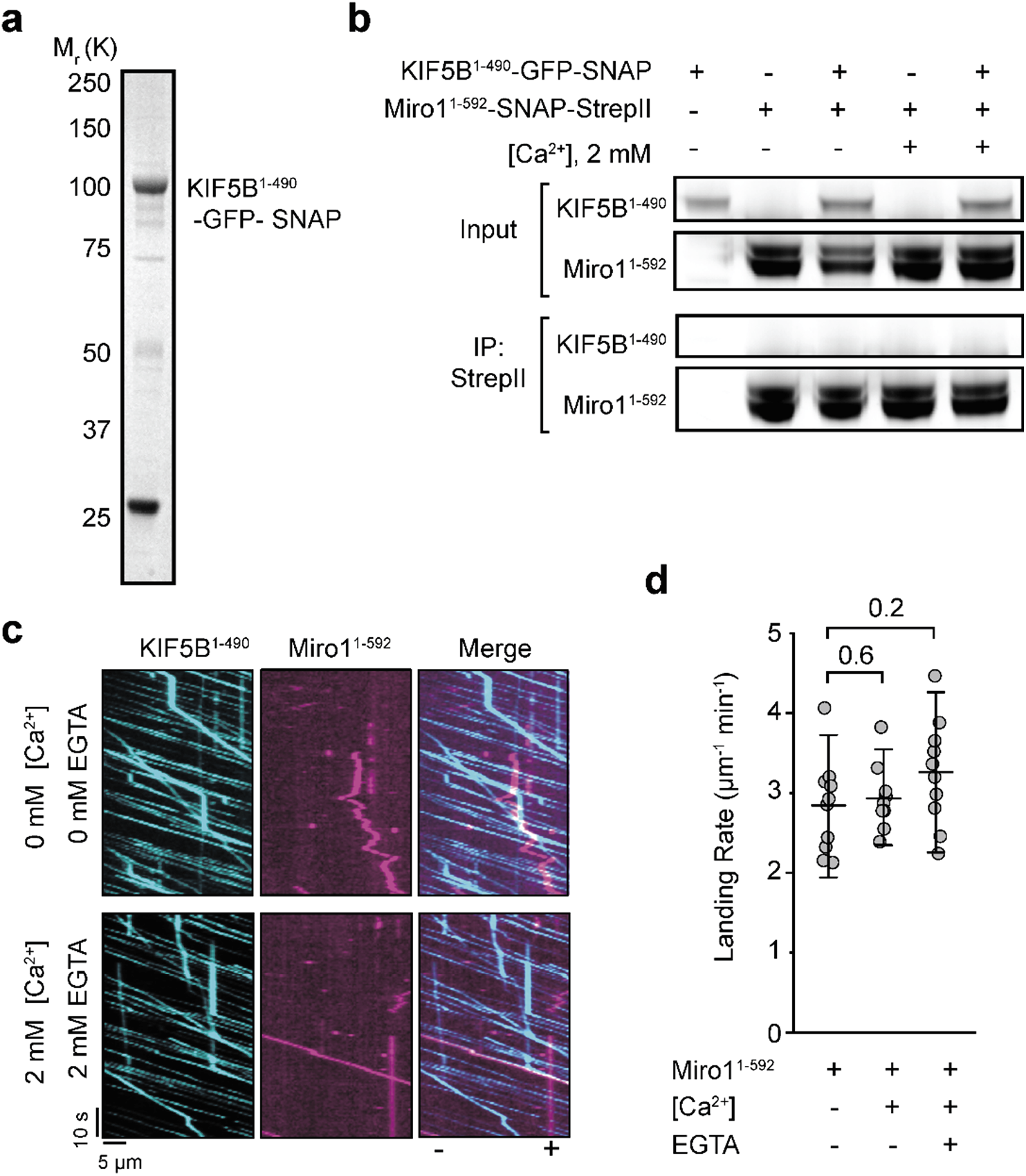
Miro1 does not interact with the kinesin motor domain. **a**, Denaturing gel picture of purified KIF5B^1-490^-GFP-SNAP. **b**, In vitro immunoprecipitation shows no interaction between purified KIF5B^1-490^ and Miro1^1-592^ in the presence and absence of 2 mM Ca^2+^. **c**, Representative two-color kymographs of KIF5B^1-490^ and Miro1^1-592^ in the presence and absence of 2 mM Ca^2+^ and 2 mM EGTA. Assays were performed in the absence of TRAK. **d**, The landing rate of KIF5B^1-490^ in the presence and absence of 2 mM Ca^2+^ and 2 mM EGTA. The center lines and whiskers represent the mean and s.d., respectively (*n* = 10 MTs for all conditions, three independent trials). P-values are calculated from a two-tailed t-test.

**Extended Data Fig. 7.**
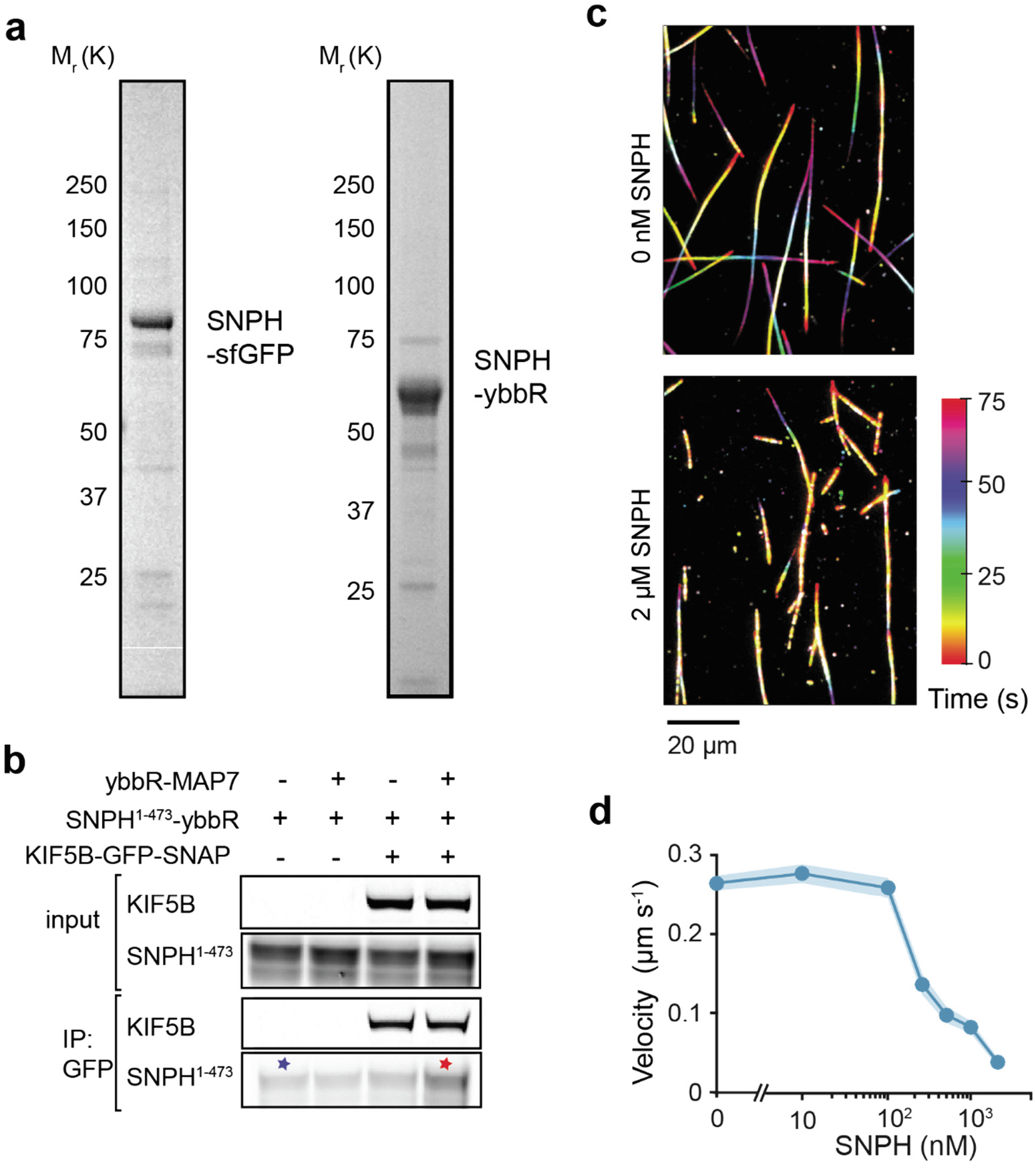
Effect of SNPH on the MT gliding by constitutively-active kinesin. **a**, Denaturing gel pictures of purified SNPH^1-473^-sfGFP and SNPH^1-473^-ybbR. **b**, In vitro immunoprecipitation shows no interaction between purified kinesin (KIF5B) and SNPH^1-473^ above background (blue star) and a minor interaction in the presence of MAP7 (red star). **c**, Representative color-coded projections of gliding MTs driven by 2.5 nM constitutively-active KIF5B^1-560^-GFP motors in the presence or absence of 2 μM SNPH-sfGFP. **d**, MT gliding velocity driven by KIF5B^1-560^ motors in the presence of different SNPH^1-473^-sfGFP concentrations (mean ± s.e.m., *n* = 31, 53, 67, 51, 71, 52, 68 MTs from left to right, three independent trials).

## Video Legends

**Supplementary Video 1: Processive motility of individual DDT complexes on MTs.** One-color imaging of DDT complexes assembled with unlabeled dynein and dynactin, and LD555-labeled TRAK adaptors on surface-immobilized MTs in the presence or absence of 1 μM Lis1. Images were acquired at 250 ms per frame.

**Supplementary Video 2: Processive motility of individual KT complexes on MTs.** Two-color imaging of LD655-kinesin (cyan) and LD555-TRAK adaptors (red) on surface-immobilized MTs in the presence or absence of 50 nM MAP7. Images were acquired at 250 ms exposure time per frame.

**Supplementary Video 3: Live imaging of DDKT_1_^1-400^ complex motility on MTs.** Three-color imaging of Alexa488-kinesin (KIF5B, cyan), LD555-TRAK1^1-400^, and LD655-dynein (magenta) on surface-immobilized MTs in the presence of unlabeled dynactin and 5 nM MAP7. Plus-end-directed motility of dynein-dynactin when it colocalizes with kinesin is highlighted by white arrows. Images were acquired at 250 ms exposure time per frame.

**Supplementary Video 4: Motility of KM complexes are disrupted by Ca^2+^.** Two-color imaging of LD655-kinesin (KIF5B, cyan) and LD555-Miro1^1-592^ (magenta) with or without 2 mM Ca^2+^. Images were acquired at 250 ms exposure time per frame.

**Supplementary Video 5: MAP7 increases the run frequency of KM complexes on MTs.** Two-color imaging of LD655-kinesin (KIF5B, cyan) and LD555-Miro1^1-592^ (magenta) on surface-immobilized MTs in 10 nM MAP7 with or without 2 mM Ca^2+^. Images were acquired at 250 ms exposure time per frame.

**Supplementary Video 6: Motility of KTM complexes assembled with TRAK1 adaptors is unaffected by Ca^2+^.** Three-color imaging of Alexa488-kinesin (KIF5B, cyan), LD555-TRAK1 adaptors (red), and LD655-Miro1^1-592^ (magenta) on surface-immobilized MTs with (right) or without (left and middle) 2 mM Ca^2+^. Assays were conducted in the presence of 10 nM MAP7. Images were acquired at 250 ms exposure time per frame.

**Supplementary Video 7: Motility of KTM complexes was not observed when TRAK1 was replaced with TRAK2.** Three-color imaging of Alexa488-kinesin (KIF5B, cyan), LD555-TRAK2 adaptors (red), and LD655-Miro1^1-592^ (magenta) on surface-immobilized MTs with (right) or without (left and middle) 2 mM Ca^2+^. Assays were conducted in the presence of 10 nM MAP7. Images were acquired at 250 ms exposure time per frame.

**Supplementary Video 8: SNPH reduces the MT landing rate and velocity of kinesin.** Motility of LD655-kinesin (cyan) on surface-immobilized MTs in the presence or absence of 500 nM GFP-SNPH (red). SNPH decorates the MT surface. Images were acquired at 250 ms exposure time per frame.

**Supplementary Video 9: SNPH inhibits MT gliding by kinesin.** Gliding motility of Cy5-MTs on surfaces decorated with 2.5 nM kinesin-GFP in the presence or absence of 500 nM GFP-SNPH. Images were acquired at 150 ms exposure time per frame.

## Raw Gel Images

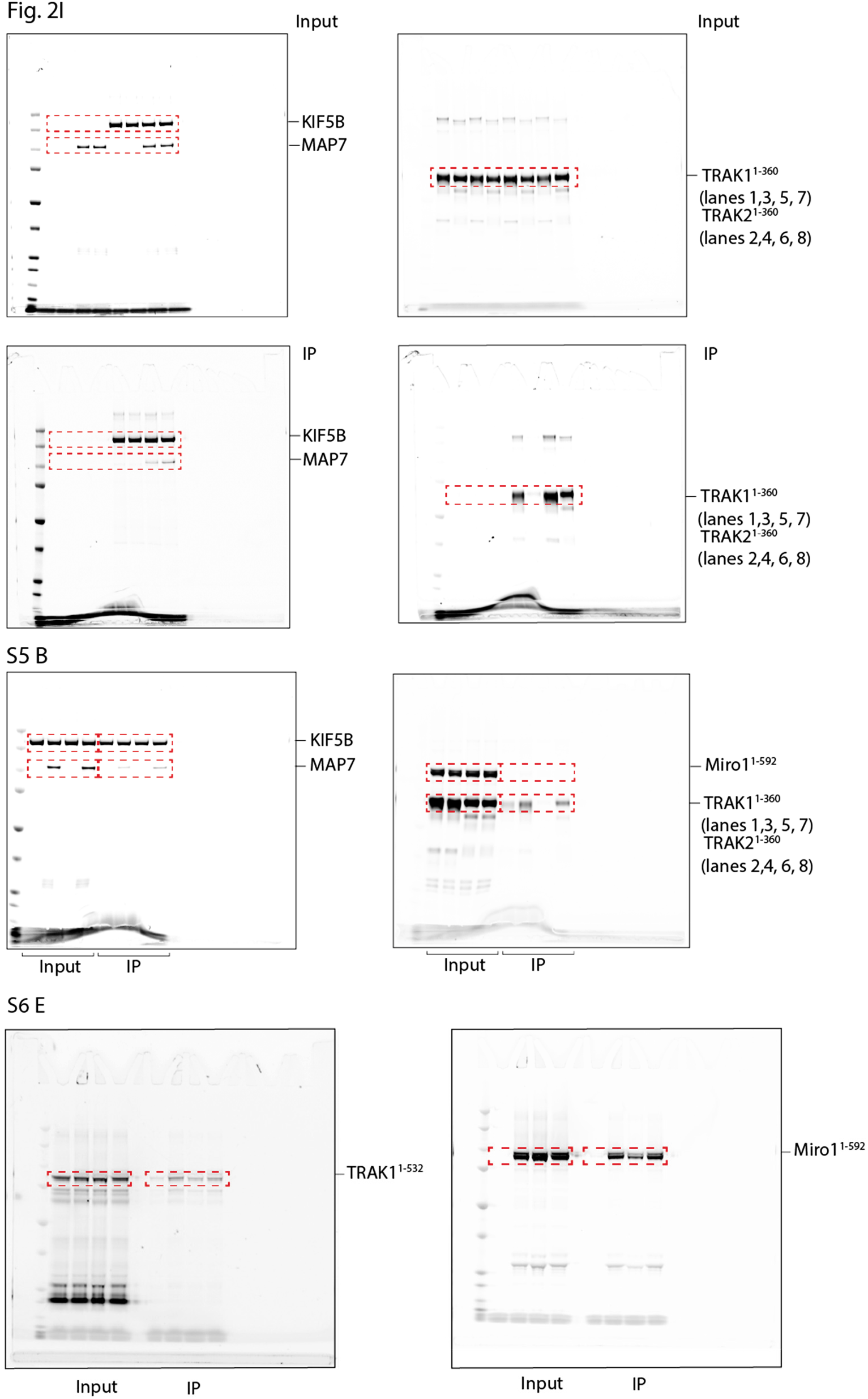

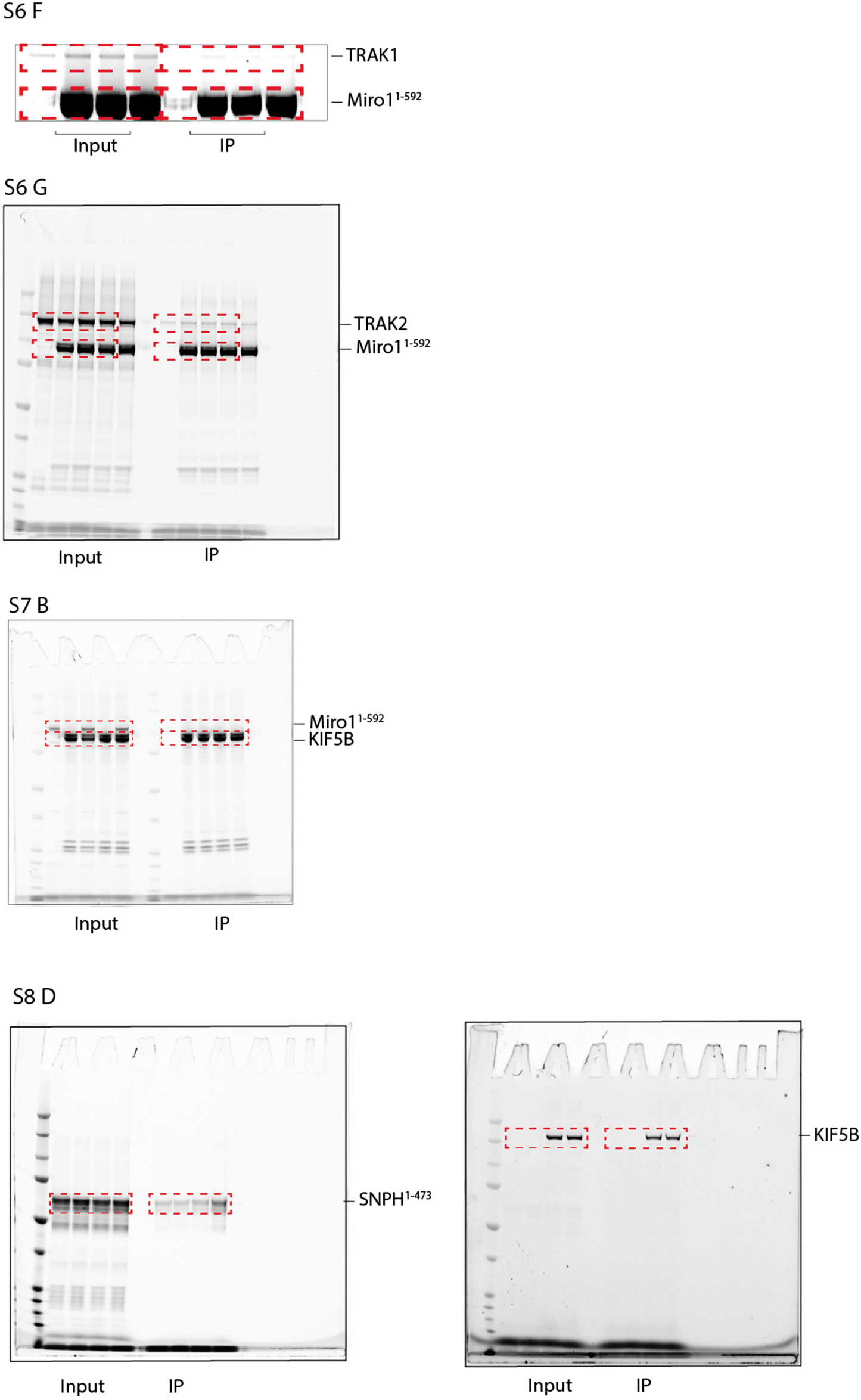

